# Establishment of a *Mycoplasma hyorhinis* challenge model in five-week-old piglets

**DOI:** 10.1101/2023.01.17.524379

**Authors:** Dorottya Földi, Zsófia Eszter Nagy, Nikolett Belecz, Levente Szeredi, József Földi, Anna Kollár, Miklós Tenk, Zsuzsa Kreizinger, Miklós Gyuranecz

## Abstract

*Mycoplasma hyorhinis* is an emerging swine pathogen bacterium with high prevalence worldwide. The main lesions caused are arthritis and polyserositis and the clinical manifestation of the disease may result in significant economic losses due to the decreased weight gain and enhanced medical costs.

Our aim was to compare two challenge routes to induce *M. hyorhinis* infection using the same clinical isolate. Five-week-old, Choice hybrid pigs were inoculated on two consecutive days by intravenous route (Group IV-IV) or by intravenous and intraperitoneal route (Group IV-IP). Mock infected animals were used as control (Control Group). After challenge, the clinical signs were recorded for 28 days, after which the animals were euthanized. Gross pathological and histopathological examinations, PCR detection, isolation and genotyping of the re-isolated *Mycoplasma* sp. and culture of bacteria other than *Mycoplasma* sp. were carried out. ELISA test was used to detect anti-*M. hyorhinis* immunoglobulins in the sera of all animals. Pericarditis and polyarthritis were observed in both challenge groups, however the serositis was more severe in Group IV-IV. Statistically significant differences were detected between the challenged groups and the control group regarding the average daily weight gain, pathological scores and ELISA titres. Additionally, histopathological scores in Group IV-IV differed significantly from the scores in the Control Group. All re-isolated strains were the same or a close genetic variant of the original challenge strain. Our results indicate that both challenge routes are suitable for modelling the disease. However, due to the more severe pathological lesions and the more natural-like route of infection in Group IV-IV, the two-dose intravenous challenge is recommended by the authors to induce serositis and arthritis associated with *M. hyorhinis* infection.

## Introduction

*Mycoplasma hyorhinis* is an emerging pathogenic bacterium of swine, distributed worldwide with an estimated prevalence of 50-70% in the herds (Pieters & Maes, 2019; Roos et al., 2019). *M. hyorhinis* colonises the upper respiratory tract and tonsil of sows, which are asymptomatic carriers of the bacterium. Piglets get infected directly from the nasal secretions of sows, and later from each other, especially after weaning (Clavijo et al., 2017). Clinical signs usually appear between three to ten weeks of age. Although the susceptibility to the infection decreases after this age, pigs can get infected even up to 16 weeks of age (Martinson et al., 2017). The pathomechanism of systemic spread is still not fully understood. Predisposing factors, such as inadequate housing conditions or weaning and decreasing maternal antibodies around three weeks of age can all contribute to the disease (Clavijo et al., 2019).

The first clinical signs appear on the third to tenth days after exposure and include fever and lethargy (Gomes Neto et al., 2012). Later coughing, laboured breathing and dyspnoea can appear due to serofibrinous pleuritis, pericarditis and peritonitis. Additionally, arthritis with swollen joints and lameness can be observed in pigs (Barden & Decker, 1971). Rarely, *M. hyorhinis* infection can cause otitis (Morita et al., 1995), conjunctivitis (Resende et al., 2019) and meningitis (Bünger et al., 2020). Affected pigs show growth retardation which can be evident even five months after infection (Barden & Decker, 1971). As a secondary pathogen *M. hyorhinis* can aggravate the clinical signs of other infections like porcine circovirus 2 associated diseases, porcine respiratory disease complex and enzootic pneumonia (Pieters & Maes, 2019). Decreased weight gain and cost of medical treatments result in significant economic losses. As no commercial vaccine is available in Europe, prevention mainly relies on decreasing predisposing factors, however metaphylactic antibiotic treatment is often required. There are some *M. hyorhinis* challenge models in the literature suggesting different inoculation routes. Not all of these models are suitable for vaccine efficacy studies as with some of the suggested challenge routes not all typical lesions can be induced (Martinson, Minion, et al., 2018).

Our aim was to compare the effects of experimental infections of two distinct inoculation routes with the same virulent *M. hyorhinis* strain by studying the colonisation of the bacteria, clinical signs, immune response and macroscopic and microscopic alterations. Accordingly, the examinations also aimed to establish a challenge model for future vaccine efficacy studies.

## Materials and methods

### Challenge material

The *M. hyorhinis* isolate used during this study was isolated from the pericardium of an affected pig originated from Hungary in 2016. The initial isolation was carried out using the filter cloning technique in Mycoplasma Experience Medium (Mycoplasma Experience Ltd., Bletchingley, UK). The challenge material was prepared freshly for each challenge day by inoculating the isolate 48 hours prior challenge and incubating at 37°C. The copy number determination was carried out on the day of the challenge. The number of colour changing units (CCU/ml) were calculated by plate micro-dilution from the highest dilution showing colour change (red to yellow shift) (Hannan, 2000).

### Experimental animals

Sixteen, four-week-old Choice hybrid piglets were transported to the animal house of the Veterinary Medical Research Institute six days prior infection. The animals were obtained from a farm with low *M. hyorhinis* prevalence and high health status (free from: brucellosis, leptospirosis, Aujeszky’s disease, porcine reproductive and respiratory syndrome, swine dysentery, atrophic rhinitis, *Actinobacillus pleuropneumoniae, Mycoplasma hyopneumoniae*, lice and mange). The *M. hyorhinis* free status of the piglets were checked before challenge by real-time PCR testing and *Mycoplasma* culture of nasal swabs.

Upon arrival, the animals were weighed and randomly divided into three groups with similar average weight. The groups were housed in separate pens, feed and water were provided *ad libitum*. The experiment was approved by the National Scientific Ethical Committee on Animal Experimentation under reference number: PE/EA/746-7/2021.

### Challenge routes

Group IV-IV (n=6) was inoculated by intravenous (IV) route on days 0 and 1 (D0, D1) with 10 ml 10^6^ CCU/ml challenge material. Group IV-IP (n=6) was challenged IV on D0 with 10 ml 10^6^ CCU/ml challenge material and intraperitoneal (IP) route on D1 with 20 ml 10^6^ CCU/ml challenge material. Total challenge dose was 2×10^7^ CCU/pig and 3×10^7^ CCU/pig in Group IV-IV and IV-IP, respectively. The controls (n=4) were inoculated by IV route on D0. Two of these animals were inoculated by IV route and the remaining two pigs by IP route on D1. Animals in the Control Group received only sterile liquid media in the same volume as the challenged groups.

### Clinical observation

The animals were observed daily from D0 until the end of the study at D28. Clinical signs of arthritis (swollen joints, lameness) and respiratory disease (coughing or laboured breath) were recorded. Body temperatures were measured daily from D-2. Body weight measurement, blood and nasal swab sampling were carried out twice a week. Schedule of events are summarized in Table 1. Average daily weight gain (ADWG) was calculated by subtracting the weight measured at D-6 from the weight measured at D27 and dividing it by the number of days past (n=33).

**Table 1:** Schedule of events, challenge routes and doses. ^†^IV-intravenous, IP-intraperitoneal, CCU-colour changing unit

| Time | Event<sup>†</sup> | Challenge dose<sup>†</sup> |
| --- | --- | --- |
| D-6 | Arrival of 16 four-week-old piglets<br>Body weight measurement |  |
| D-2 | Blood sampling<br>Collection of nasal swabs for PCR and *Mycoplasma* isolation<br>Body temperature measurement |  |
| D-1 | Body temperature measurement |  |
| D0 | IV challenge of all groups | 10 ml 10<sup>6</sup> CCU/ml challenge material (Groups IV-IV and IV-IP) or 10 ml sterile broth (Control Group) |
| D1 | IV challenge of Group IV-IV and two animals from Control Group<br>IP challenge of Group IV-IP and two animals from Control Group 3 | IV: 10 ml 10<sup>6</sup> CCU/ml challenge material (Group IV-IV) or 10 ml sterile broth (Control Group)<br>IP: 20 ml 10<sup>6</sup> CCU/ml challenge material (Group IV-IP) or 20 ml sterile broth (Control Group) |
| D0-D27 | Daily body temperature measurement and clinical observations<br>Twice a week body weight measurement<br>Collection of nasal swabs for PCR and *Mycoplasma* isolation<br>Blood sampling |  |
| D28 | Euthanasia<br>Pathological examination<br>Sample collection for PCR, histopathology and bacteriology |  |

### Isolation, DNA extraction and PCR

Nasal swabs for *Mycoplasma* isolation and PCR were taken twice a week from all animals throughout the study. Separate swab samples for *Mycoplasma* isolation and PCR were collected during necropsy as well (see below). For *Mycoplasma* isolation swabs were cut into Mycoplasma liquid media (Mycoplasma Experience Ltd.), washed then filtered by 0.45 µm pore size filters and incubated at 37°C until colour change.

DNA extraction from the swabs and colour changed broths were performed by ReliaPrep gDNA Tissue Miniprep System (Promega Inc., Madison, USA) according to the manufacturers’ instructions. For the *M. hyorhinis* species specific real-time PCR, previously published (Resende et al., 2019) primers targeting the 16S rRNA gene were optimized. Primer and probe sequences were the following: Forward primer 5’-CGT ACC TAA CCT ACC TTT AAG -3’, Reverse primer 5’-TAA TGT TCC GCA CCC C -3’, Probe 5’-FAM-CCG GAT ATA GTT ATT TAT CGC ATG ATG AG-BHQ -3’. The PCR was performed using a Bio-Rad C1000 Touch™ Thermal Cycler, CFX96™ Real-Time System (Bio-Rad Laboratories Inc., USA). The PCR master mix consisted of 6 µl 2× qPCRBIO Probe Mix No-ROX (PCR Biosystems Ltd., UK), 0.4 µl of each primer (10 µM), 0.2 µl probe and 2 µl DNA in the final volume of 12 µl. PCR conditions were the following 95°C for two minutes, 45 cycles of 95°C for 5 seconds and 60°C for 20 seconds. In order to test the sensitivity of the developed assays, tenfold dilutions of the type strain (NCTC 10130) were used in the range of 10^6^-10^0^ copy number/µl. Template copy number was calculated with the help of an online tool (http://cels.uri.edu/gsc/cndna.html) by measuring the concentration of DNA of pure *M. hyorhinis* culture by Nanodrop 2000 Spectrophotometer (Thermo Fisher Scientific Inc., USA). The lowest DNA concentration giving specific signal was considered the detection limit of the assay. The specificity was tested by including *M. hyopneumoniae, M. hyosynoviae* and *M. flocculare* in the analyses.

Necropsy samples were also tested for the presence of *M. hyopneumoniae* (Wu et al., 2019) and *M. hyosynoviae* (Martinson, Minion, et al., 2018) by PCR. *M. hyorhinis* positive isolates were genetically characterized by multi-locus sequence typing (MLST: costly and robust genotyping system) and multiple-locus variable-number tandem-repeat analysis (MLVA: fast and cheap genotyping system with high-resolution) by previously published assays (Földi et al., 2020).

### Gross pathological examination

Joints of carpus, elbow, tarsus and stifle on both sides were opened and examined for the signs of arthritis. The thoracic and abdominal cavity (pleura, pericardium, peritoneum) were checked for serositis. Body condition, skin, subcutaneous tissues, musculoskeletal system, eyes and conjunctiva, nasal, and oral cavity, trachea, lungs, heart, lymph nodes, gastrointestinal system, liver, spleen, kidney and brain were also checked for lesions. Scoring system of the gross pathological examination is detailed in Supplementary table 1. Lesions of joints and serosa were scored to reflect severity based on previously described criteria (Martinson, Zoghby, et al., 2018). Total scores were calculated by summarizing all organ scores.

Swab samples for bacterial culture, *M. hyorhinis* isolation and PCR were taken from the conjunctiva, lung, serosa, the four examined joints and brain. Joints on both sides were sampled with the same swab.

### Histological examination

Samples for histopathology were collected from conjunctiva, choana, tonsilla, trachea, lungs (7 lobes), pericardium, heart, mediastinal and mesenterial lymph nodes, liver, spleen, kidney, joints and brain (cerebrum, cerebellum, brain stem). Tissue samples were fixed in 10% formaldehyde, embedded in paraffin then 4 µm thick sections were cut and stained with haematoxylin and eosin (H&E) and examined by light microscope. Data about the scoring system of *M. hyorhinis* infected tissue lesions are scarce in the literature. Given the limited number of examined animals in the present study, the establishment of a general scoring system was not possible either. Therefore, lesions were categorized based on the comparison of the severity of the histopathological changes to each other, and scores were assigned as follows: 0-no lesion, 1-mild lesion, 2-moderate lesion, 3-severe lesion in the given organ.

### Bacteriology

Presence of bacterial pathogens other than *Mycoplasma* sp. were tested by culturing the necropsy samples on Columbia sheep blood agars (Biolab Inc., Hungary) and sheep blood agars supplemented with nicotinamide adenine dinucleotide (Sigma-Aldrich Co., USA) at the final concentration of 20 µg/ml. The agar plates were incubated at the presence of 5% CO_2_ at 37°C for 48 hours.

### Serology

Sera were tested in duplicates by an in-house ELISA, using an antigen prepared according to the sarcosyl assay previously described for *M. gallisepticum* (Stipkovits et al., 1993). Briefly, to prepare the antigen, six clinical isolates of *M. hyorhinis* were propagated (Supplementary table 2). After colour change the isolates were mixed, washed and treated with 0.5% sarcosyl. Protein content of the antigen was determined with Coomassie (Bradford) Protein assay kit (Thermo Fisher Scientific Inc.) according to the manufacturer’s instructions.

96-well ELISA plates were coated with the antigen diluted to the concentration of 1.25 µg/ml in phosphate buffered saline (PBS, pH 7.4). After blocking with 1% gelatine from cold water fish skin (Sigma-Aldrich Co.) each well was incubated with serum sample diluted to 1:100 in PBS, followed by a horseradish peroxidase conjugated rabbit anti-swine immunoglobulin (Dako A/S, Denmark) diluted to 0.125 µg/ml in PBS. The reaction was visualized with tetramethylbenzidine (TMB, Diavet Ltd., Hungary) substrate and the optical density of the solution was measured at 450 nm using a Multiscan FC reader (Thermo Fisher Scientific Inc.). Blood samples were centrifuged after collection and the sera were kept at −70°C. Each serum sample were thawed only once. Each plate contained a negative control (mix of the sera of each Control animal taken at D28 from this study) a positive control (mix of the sera of each animal in Group IV-IV taken at D28 from this study) and a background control, where PBS was measured instead of the serum sample. The mean OD value of the background control was subtracted from the mean OD values of the samples and the controls (Terato et al., 2017). The assay was considered valid if the negative to positive ratio of corrected OD values were under 40%. The sample to positive ratios (S/P%) were calculated and the sample was considered positive when S/P%>40% (Merodio et al., 2021).

### Statistical analyses

Statistical analyses were accomplished with R programme (R Core Team, 2021). To compare the effect of the different challenge routes statistical analysis of the ADWG, pathological scores (separately for the joints, serosa of pericardium, pleura and peritoneum and summary of scores), histopathological scores (separately for the joints, serosa of pericardium, pleura and peritoneum and summary of scores) and ELISA results from the last sampling were performed. In case of the pathological and histopathological scores first a Kruskal-Wallis non-parametric ANOVA test was carried out to determine whether the difference among the medians of the three study groups are statistically significant or not. If the results of the Kruskal-Wallis test were significant a Dunn’s test was performed to determine exactly which groups are different by making pairwise comparisons between each group. Since multiple groups were considered at the same time, p-values were adjusted for multiple comparisons by Bonferroni method. In case of the ADWG and the ELISA results instead of the non-parametric test a one-way ANOVA followed by Tukey multiple comparisons of means was performed after the normal distribution of the data was tested by Shapiro-Wilk normality test.

## Results

### Clinical observations

No clinical alterations were detected in the Control Group throughout the study. No body temperature higher than 40.3°C was recorded during the study. One pig in Group IV-IP had body temperatures higher than 40°C on three consecutive days (D4-6, Supplementary table 3). No respiratory signs were recorded in the challenge groups.

Swollen joints were detected as early as D6 in Group IV-IP and D8 in Group IV-IV. Typically, the first swollen joint was one of the tarsal joints. By D15 all pigs in Group IV-IP had at least one swollen joint, 3/6 pigs had two swollen tarsi and in one animal joints of the front legs were also affected. In Group IV-IV swollen tarsal joint was observed in 4/6 pigs (one side only) and in one animal both tarsi were affected, while no swollen joints were detected in one pig (Supplementary table 4).

Weight gain dynamics of the different groups are shown in Figure 1 and detailed in Supplementary table 5. Average starting weight of the groups were 10.5 kg (SD 1.2), 10.3 kg (SD 1.2) and 10.3 kg (SD 1.1) in Group IV-IV, IV-IP and Control while at the end of the study average body weight of the groups were 18.0 kg, 16.0 kg and 22.0 kg respectively. Mean ADWG was 223 g, 170 g and 350 g in Group IV-IV, IV-IP and Control Group. Significant differences in ADWG were detected between Groups Control and IV-IV (p=0.05) and Groups Control and IV-IP (p<0.01; Supplementary data 1).

**Figure 1:**
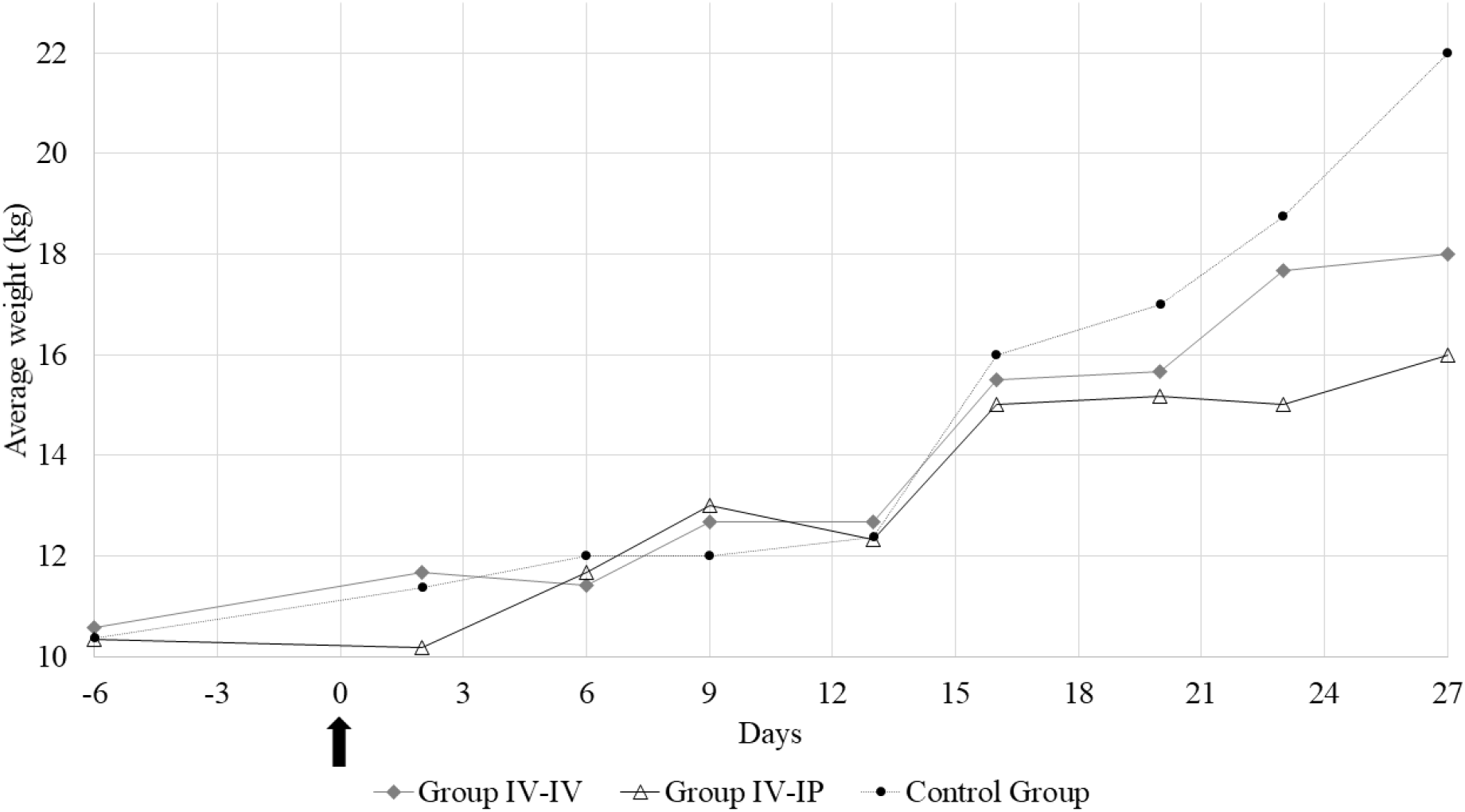
Average weight of the study groups at each sampling point. The arrow marks the first day of the challenge.

### Mycoplasma isolation, PCR and bacteriology

Sensitivity of the reaction with the optimized primers were 10^1^ copies/reaction, no cross reactions were detected for *M. hyopneumoniae, M. hyosynoviae* and *M. flocculare*. Nasal swabs of all animals were negative for *M. hyorhinis* by PCR and isolation at the beginning of the study (D-2). After the inoculation of the pigs one sample from each challenged group was positive by isolation which were positive also by PCR either at same time or at different sampling times. These animals remained PCR positive for two-four consecutive sampling points. Further two animals in Group IV-IV and one animal in Group IV-IP were PCR positive as well at one sampling point. All nasal samples of Control Group were negative by PCR and isolation for *M. hyorhinis* throughout the study (Supplementary table 6).

Samples collected from the conjunctiva and meninx during necropsy were negative for the tested mycoplasmas in all animals, while one lung sample in Group IV-IV was positive for *M. hyorhinis* by PCR. Three samples from different serosa (pleura, pericardium, peritoneum) were positive by PCR as well in Group IV-IV. High number of joint samples were positive by PCR in both challenged groups. In Group IV-IV 2/6 stifle, 4/6 elbow, 5/6 tarsus and 4/6 carpus samples were positive for *M. hyorhinis* by PCR. While in Group IV-IP 2/6 stifle, 4/6 elbow, 4/6 tarsus and 1/6 carpus samples were positive. All samples from Control Group were negative for *M. hyorhinis* (Supplementary table 7). *M. hyopneumoniae* or *M. hyosynoviae* were not detected in any samples collected during necropsy.

During the challenge study two nasal isolates and isolates from six necropsy samples (tarsal, carpal, elbow and stifle joints) were collected and their genotypes were first determined by MLVA. Two re-isolates in Group IV-IP differed from the challenge strain on one allele (MHR444; Supplementary table 8). They were microvariants due to within host evolution. The sequence types of these two isolates, two other isolates from the same animals and one isolate from Group IV-IV were also determined by MLST. All the re-isolated strains showed the same sequence type (ST) with MLST as the challenge strain (Supplementary data 2). MLST and MLVA trees are shown in Supplementary figure 1.

None of the cultures of the necropsy samples showed growth of pathogenic bacteria that could also be associated with the lesions, other than *M. hyorhinis*.

### Gross pathological examination

Arthritis of at least one joint was observed in all pigs in the challenge groups. Mild to severe arthritis was found in all joints examined in one pig and in three joints examined in another pig in Group IV-IV. A single joint was affected in the remaining four animals in this group. Mild to severe arthritis was found in three, two or one joints of two-two pigs in Group IV-IP. The arthritis manifested as serous or purulent inflammation (Figure 2A) and was detected most often in the tarsus (8/12) followed by elbow (6/12), stifle (5/12) and the carpus (4/12) on one or on both sides. In addition to arthritis, erosions in the cartilage were evident in the tarsal joint of two animals in Group IV-IP, which indicates a prolonged time of inflammation of the joint (Figure 2B).

**Figure 2:**
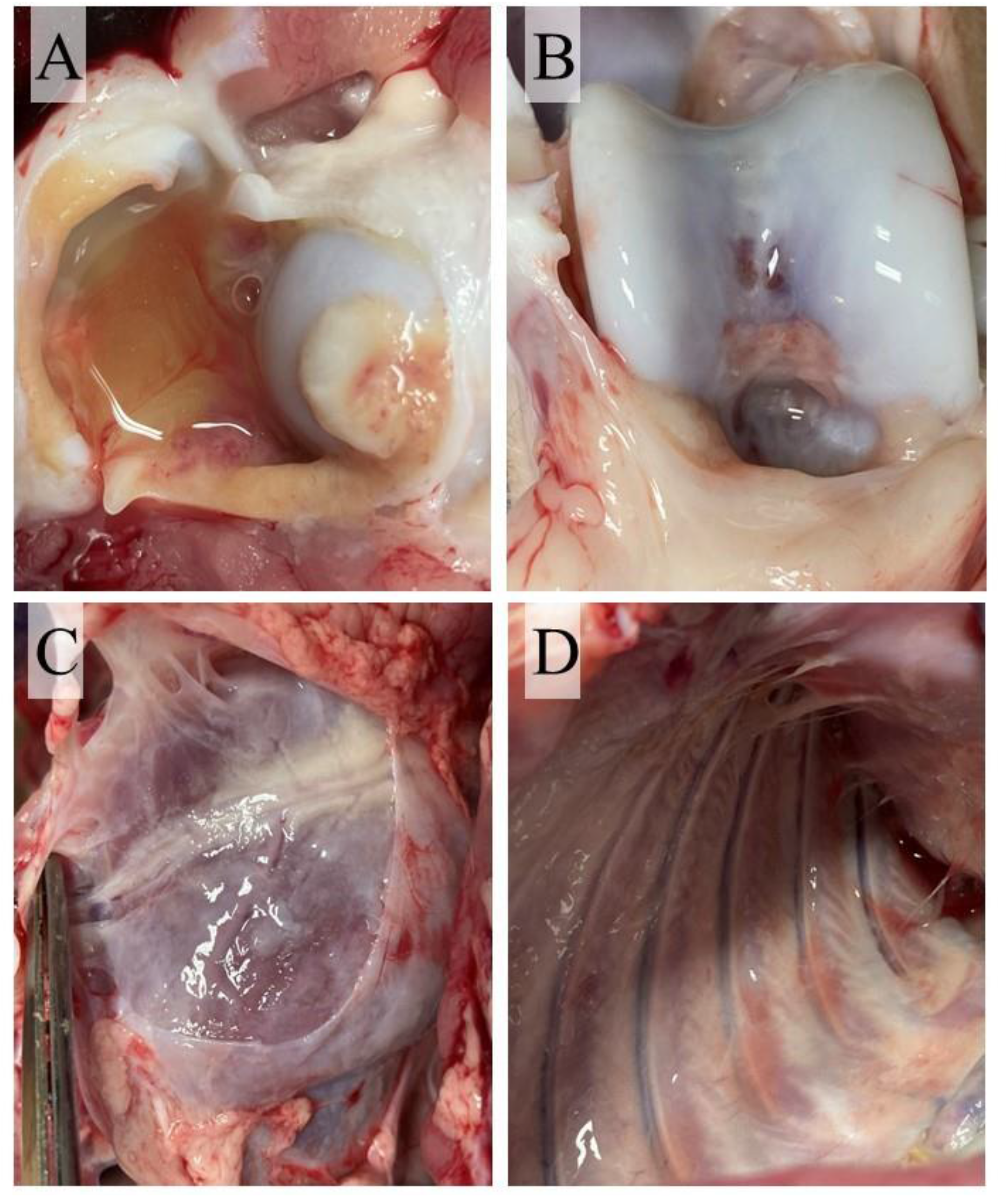
Typical lesions of *Mycoplasma hyorhinis* infection. A-Joint with excess synovial fluid. B-Cartilage erosion. C-Serofibrinous pericarditis. D-Serofibrinous pleuritis.

Diffuse, severe, chronic pericarditis presenting large amount of connective tissue was detected in two animals in both groups (Figure 2C). Additionally, mild or moderate chronic pleuritis presenting filaments of connective tissues were detected in two animals (Figure 2D) and mild chronic peritonitis presenting filaments of connective tissues occurred in one other animal in Group IV-IV.

Macroscopic scores of lesions in the affected organs are demonstrated in Figure 3. No gross pathological alterations were found in the remaining organs examined. No gross pathological lesions were detected in any examined organs in the Control Group. Body condition in all groups was normal. Detailed pathological scores are given in Supplementary table 1.

**Figure 3:**
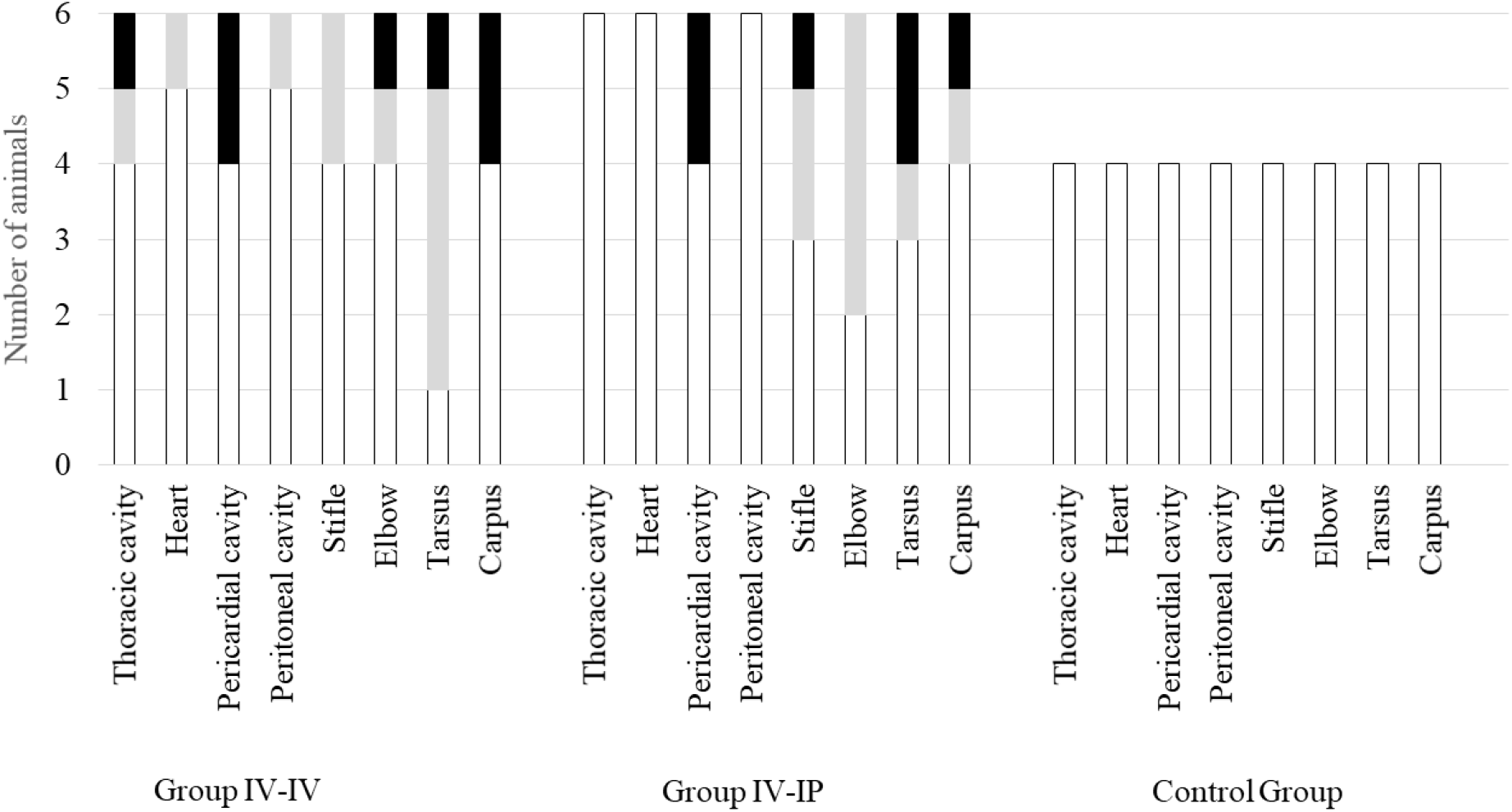
Scores of macroscopic lesions of the affected organs of the study groups. Organs were scored between 0-2 based on the severity of the lesion, except for the heart where score 1 was given in case of any lesion (Supplementary table 1). In the charts white indicates the number of animals with score 0, light grey indicates the number of animals with score 1 and black indicates the number of animals with score 2. The number of animals in each group is indicated on the Y-axis: the challenge groups consisted of six animals, while the control group involved four animals.

Significant differences in pathological scores were detected when scores of joint lesions and total scores of groups were compared: pathological scores in both challenge group differed significantly from the Control Group (p=0.03 and p=0.02 regarding joint lesions, p=0.02 and p=0.03 regarding total scores for Group IV-IV-Control and Group IV-IP-Control, respectively), but not from each other in both cases. No significant difference was found when scores of serosa lesions were compared (Supplementary data 1).

### Histological examination

The results of the histological examination are summarised in Supplementary table 9. The main alterations were detected in the joints and in the serosa of parenchymal organs in the thoracic and peritoneal cavity. Arthritis was found in 6/6 pigs in Group IV-IV and 4/6 pigs in Group IV-IP. Mild to severe lympho-histiocytic inflammation associated with the formation of lymphoid follicle around blood vessels were detected in four animals in both Groups IV-IV and IV-IP, and the same changes with the presence of multinucleated giant cells were found in two infected animals in Group IV-IV (Figure 4A). Alterations in the pleura were present in 3/6 and 4/6 infected pigs in Groups IV-IV and IV-IP, respectively. Filamentous pleural projections consisting of connective tissue with serous-purulent inflammation and multinucleated giant cells were found in one animal in Group IV-IV (arthritis with multinucleated giant cells was also detected in the latter pig). Diffuse thickening of the pleura and filamentous pleural projections consisting of connective tissue with moderate serous-purulent inflammation were evident in 1/6 animals in Group IV-IV, while filamentous pleural projections of connective tissue without thickening of the pleura was detected in one animal (Figure 4B). Also diffuse thickening of the pleura with filamentous pleural projections consisting of connective tissue with lympho-histiocytic inflammation were evident in three animals in Group IV-IP while in this group one animal only presented diffuse thickening of the pleura without inflammation (Figure 4B). The epicardium and pericardium were affected in 3/6 and 2/6 animals in Groups IV-IV and IV-IP, respectively. Diffuse thickening of the pericardium, filamentous projections consisting of connective tissue on the pericardium and serous purulent inflammation were evident in one pig in both groups, and severe proliferation of connective tissue associated with adhesion of the epi- and pericardium were detected in two and one pigs in Groups IV-IV and IV-IP, respectively (Figure 4B). Alterations of the peritoneum was found in 1/6 infected pig in both groups. Filamentous projections consisting of connective tissue were detected on the serosa of spleen and liver of the animal in Group IV-IV, and on the serosa of the spleen of the pig in Group IV-IP. Additionally, lympho-histiocytic purulent conjunctivitis was found in two cases (one animal in both infected groups), and serous-purulent rhinitis in one case in Group IV-IV. No lesions were detected in the other organs.

**Figure 4:**
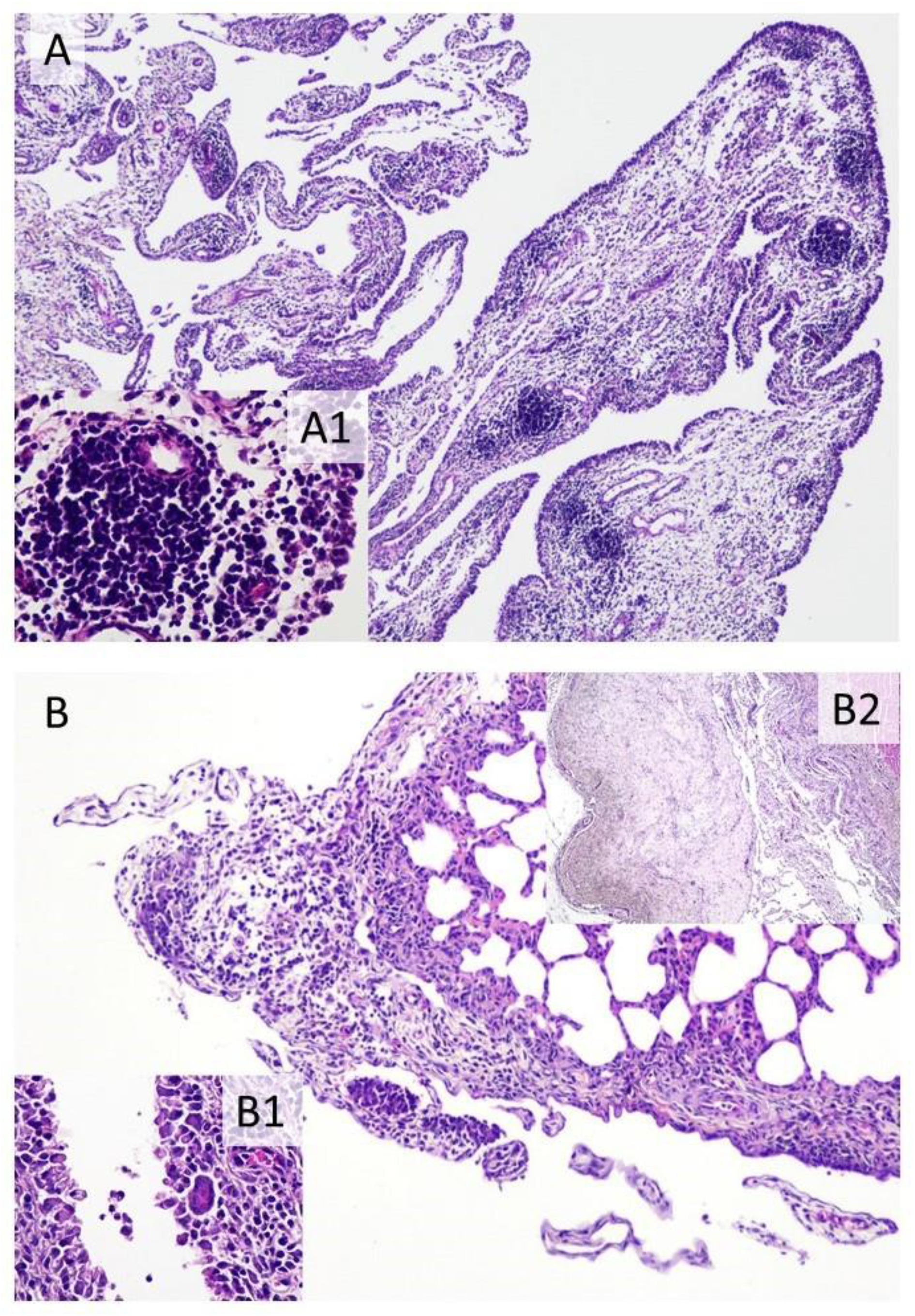
Typical histopathological changes in *Mycoplasma hyorhinis* infected piglets. A: Joint synovial membrane: Severe lympho-histiocytic inflammation associated with the formation of lymphoid follicle around blood vessels, H&E, 40×. A1: Perivascular follicle, H&E, 400×. B: Lung: Filamentous pleural projections consisting of connective tissue and serous-purulent inflammation. B1: Joint synovial membrane: Multinucleated giant cell is associated with lympho-histiocytic inflammation, H&E, 400×. B2: Pericardium: Severe proliferation of connective tissue associated with adhesion of the epi- and pericardium, H&E, 40×.

Scores of histological lesions of affected organs are shown in Figure 5. Based on the statistical analysis, scores of joints and total score differed significantly between groups. In both cases significant differences were detected between Group IV-IV and Control (p=0.04 regarding joint lesions, p=0.04 regarding total score; Supplementary data 1).

**Figure 5:**
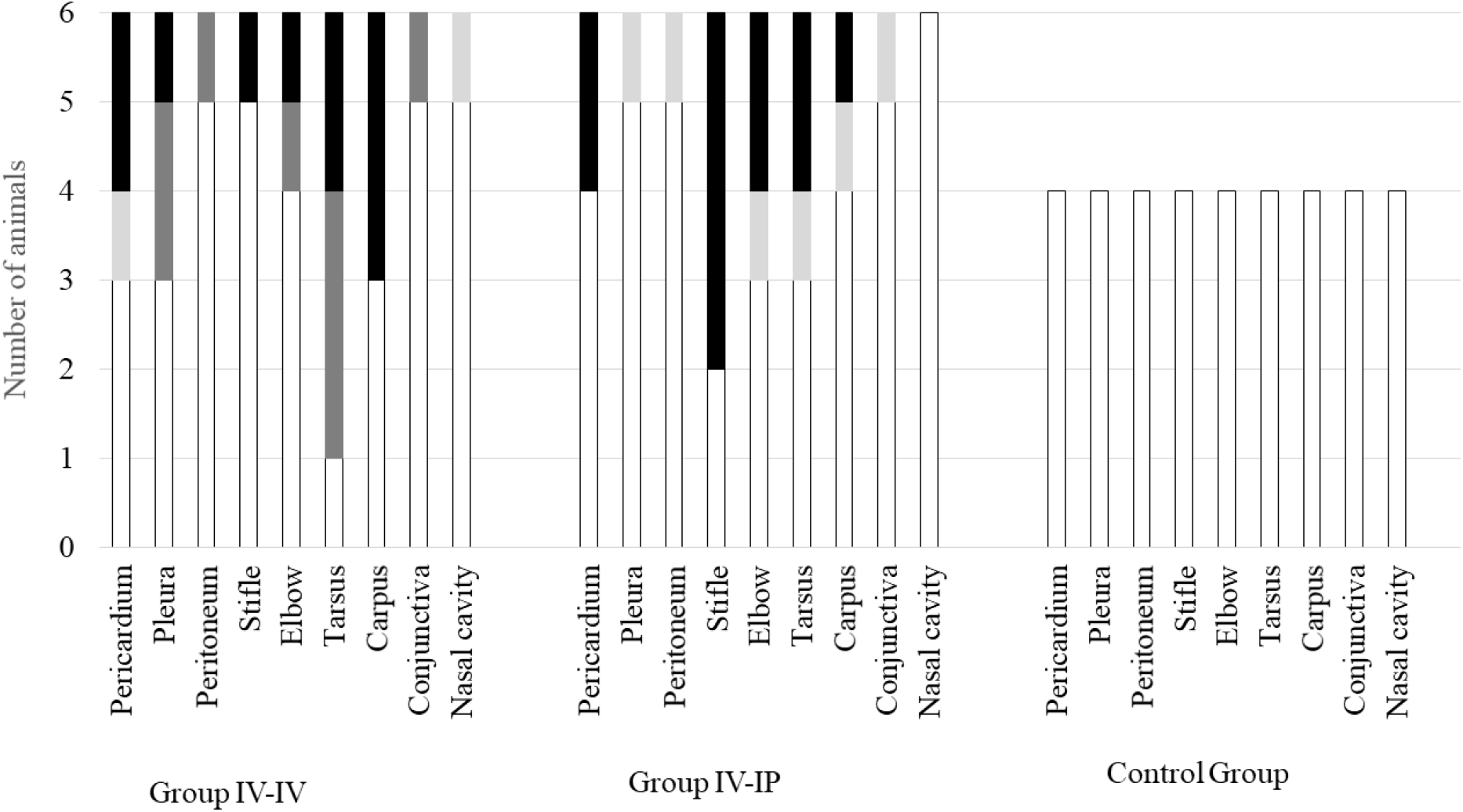
Scores of histopathologic lesions of the affected organs in the study groups. Organs were scored between 0-3 based on the severity of the lesion. In the charts white indicates the number of animals with score 0 (no lesion), light grey indicates the number of animals with score 1 (mild lesions), dark grey indicates the number of animals with score 2 (moderate lesions) and black indicates the number of animals with score 3 (severe lesions). The number of animals in each group is indicated on the Y-axis: the challenge groups consisted of six animals, while the control group involved four animals.

### Serology

All animals were serologically negative to *M. hyorhinis* at the beginning of the study. The positive serological response appeared in both challenge groups on D5, SP% of 1/6 pigs in Group IV-IV and 2/6 pigs in Group IV-IP was higher than 40%. By D28 all challenged animals were ELISA positive. However, one animal in Group IV-IP, presented positive serological response just at the last sampling point (Supplementary table 10). Mean S/P% of the groups throughout the study are demonstrated in Figure 6. Significant differences in S/P% of D28 were detected between Control and Group IV-IV (p<0.01) and Control and Group IV-IP (p<0.01; Supplementary data 1). Animals from Control Group remained negative throughout the study.

**Figure 6:**
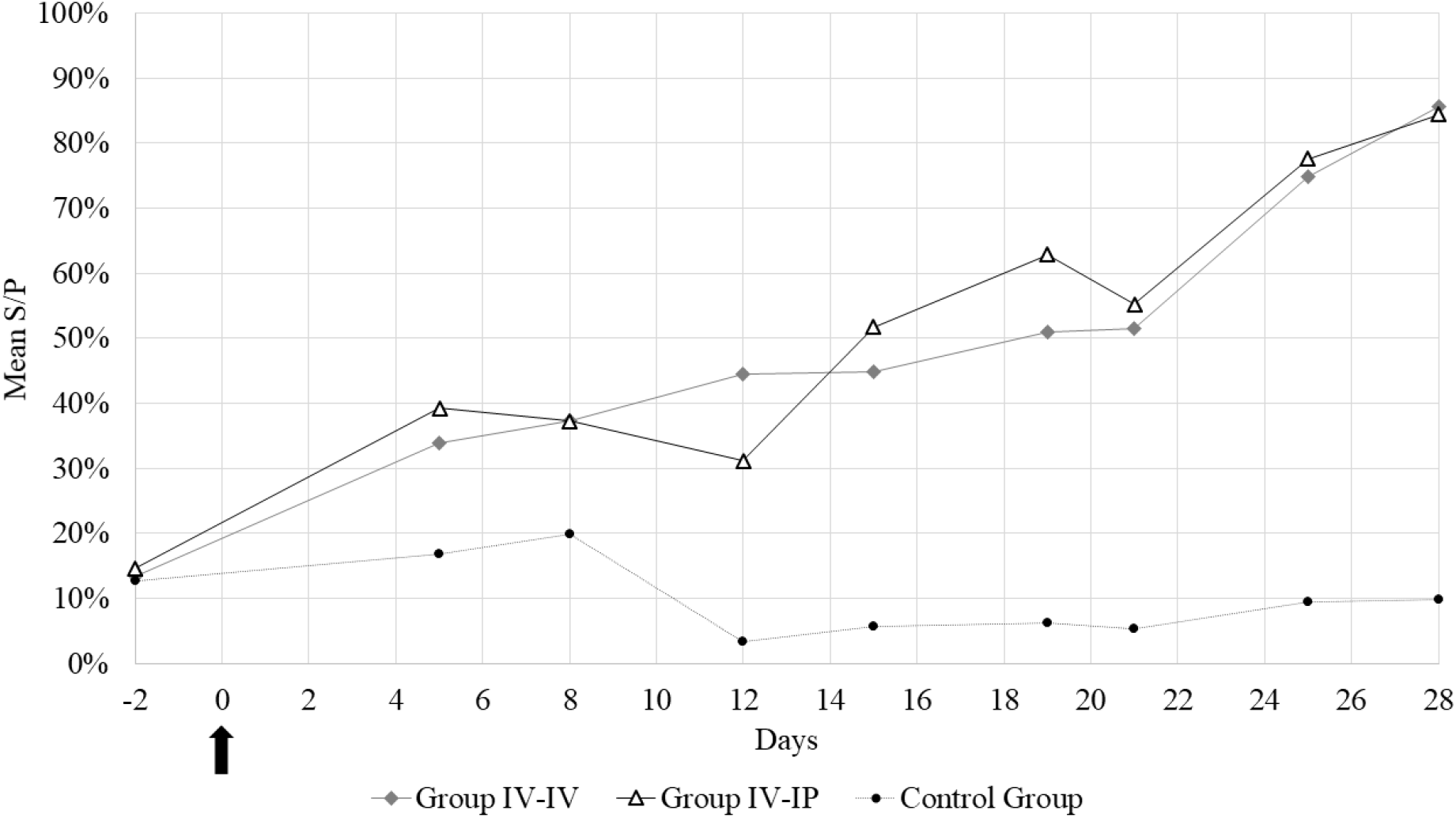
Mean sample-to-positive ratios (S/P%) of the blood samples during the study. The arrow marks the first day of the challenge.

## Discussion

Based on available literature data, the single dose intranasal or intratracheal inoculation with *M. hyorhinis* is not suitable to establish a proper challenge model as these routes usually only able to induce one aspect of the infection, mostly polyserositis and lung lesions with little or no serological conversion after infection (Fourour et al., 2019; Gomes Neto, 2014; Lee et al., 2018; Lin et al., 2006; Wei et al., 2020). Similarly, intranasal inoculation combined with tonsillar swabbing resulted in low serological conversion with no clinical signs or macroscopic lesions (Merodio et al., 2021). Time of challenge should not have an impact on the results of previous experiments as all studies used pigs at a receptive age (infected mostly at six weeks of age (Fourour et al., 2019; Gomes Neto, 2014; Lee et al., 2018; Lin et al., 2006; Merodio et al., 2021), or at ten weeks of age (Wei et al., 2020). Our study plan was based on the work of Martinson, Minion, et al. (2018) where one-dose intranasal, intravenous and intraperitoneal inoculations were compared to two- or three-dose inoculations with combined challenge routes in seven-week-old animals. The results of this study also confirmed that a single dose challenge is not sufficient to induce all typical lesions, with the mildest clinical signs observed in the intranasally infected group. On the other hand, in the intravenously infected group the rate of pigs with pericarditis and pleuritis were similar to or higher than in the groups with combined challenge routes. The authors suggested the combination of intravenous, intraperitoneal and intranasal routes on three consecutive days to induce both polyserositis and polyarthritis (Martinson, Minion, et al., 2018; Wang et al., 2022).

In the present study two challenge routes were compared by using the same virulent clinical isolate. The double dose IV challenge (which was not mentioned in previous publications) produced equal involvement of joints as the mix of IV-IP route (arthritis of at least one joint was detected in 6/6 animals in both groups), which exceeded the rate of animals affected with arthritis in the previous study (single dose IV challenge from Martinson’s work resulted arthritis in only 1/10 animal). It should be mentioned though, that with the combination of intravenous and intraperitoneal infection (Group IV-IP) clinical signs of arthritis like swollen joint and lameness appeared earlier and were more pronounced. On the other hand, in the group which was challenged by intravenous route on two consecutive days (Group IV-IV) more organs were affected by serositis than in the Group IV-IP. During necropsy, all lesions appeared chronic in both groups, therefore the reduction of the length of the study is suggested.

Although the natural route of infection is not yet fully understood, the results of the challenge models using intravenous route indicate that systemic spread of *M. hyorhinis* might happen through the circulatory or lymphatic system (Martinson, Minion, et al., 2018). This theory is further supported by the results of our study. Both of the applied challenge routes included intravenous infection, and accordingly systemic spread of *M. hyorhinis* was obtained in both cases. Furthermore, despite inoculating directly the peritoneum in Group IV-IP, peritonitis could be induced only by the double intravenous route, and overall the observed serositis was more pronounced in Group IV-IV. Nevertheless, as with both challenge methods the main lesions of *M. hyorhinis* infection were induced, both models can be recommended for the future studying of *M. hyorhinis* infection or for vaccine efficacy studies. Considering the hypothesis of the natural spread of the pathogen via the circulatory or lymphatic system and the more severe pathological lesions in Group IV-IV, two-dose intravenous challenge is recommended by the authors.

## Data availability

All data is available in the supplementary tables or supplementary data.

## Ethics Statement

The authors confirm that the ethical policies of the journal, as noted on the journal’s guidelines page, have been adhered to and the appropriate ethical review committee approval has been received. Regulations of the Hungarian Government on the use of laboratory animals were followed and our study was approved by the National Scientific Ethical Committee on Animal Experimentation under reference number: PE/EA/746-7/2021.

## Acknowledgements

The authors are thankful for Balázs Lajos for his help providing the animals meeting the study criteria.

## Supplementary information

**Supplementary data 1**

**Results of the statistical analysis**.

**Supplementary data 2**

**Aligned, concatenated sequences of the multi-locus sequence typing of the challenge strain and the re-isolates**.

**Supplementary table 1**

**Pathological scoring system and detailed pathological scores of necropsy**.

**Supplementary table 2**

**Background information of the strains used in the enzyme-linked immunosorbent assay development**.

**Supplementary table 3**

**Rectal temperatures measured during the study**.

**Supplementary table 4**

**Timescale of appearance of swollen joints**.

**Supplementary table 5**

**Results of weekly weight measurements in kilograms**.

**Supplementary table 6**

***Mycoplasma hyorhinis* specific PCR results of the nasal samples taken during the study**.

**Supplementary table 7**

***Mycoplasma hyorhinis* specific PCR results of the samples taken during necropsy**.

**Supplementary table 8**

**Repeat numbers and source of the re-isolates from this study**

**Supplementary table 9**

**Histopathological scores of *Mycoplasma hyorhinis* infected piglets**.

**Supplementary table 10**

**ELISA results of *Mycoplasma hyorhinis* infected piglets**.

**Supplementary figure 1**

**Dendrograms of multi locus sequence typing (MLST, A) and multiple-locus variable-number tandem-repeat analysis (MLVA, B)**.

A. The MLST tree was constructed by using Maximum Likelihood method, Hasegawa-Kishino-Yano model in the MegaX software (Kumar, Stecher, Li, Knyaz, & Tamura, 2018). Gene fragments from *lepA, rpoB, rpoC, gltX, valS* and *uvrA* were used, with 1000 bootstraps (only bootstrap values >70% are presented). B. Resolution of the identical sequence type of the isolates from the present study was carried out with MLVA based on Mhr205, Mhr396, Mhr438, Mhr441, Mhr442 and Mhr444 alleles. The tree was constructed by Neighbour-Joining method.

Isolates from this study are highlighted light grey on the MLST tree. MLST sequences are available in Supplementary data 2, and tandem-repeat numbers of the isolates from this study can be found in Supplementary table 8. Data of the other isolates was previously published in Földi et al., 2020.

