## Supplementary table 1-10 for "Establishment of a *Mycoplasma hyorhinis* challenge model in five-week-old piglets"

Pathological scoring system and detailed pathological results of the study

| Organ | Score | Description | Group IV-IV | | | | | | Group IV-IP | | | | | | Control Group | | | |
|---|---|---|---|---|---|---|---|---|---|---|---|---|---|---|---|---|---|---|
| | | Animal ID | 1 | 2 | 3 | 4 | 5 | 6 | 7 | 8 | 9 | 10 | 11 | 12 | 13 | 14 | 15 | 16 |
| Eyes/conjunctiva | | | 0 | 0 | 0 | 0 | 0 | 0 | 0 | 0 | 0 | 0 | 0 | 0 | 0 | 0 | 0 | 0 |
| Nasal cavity | | | 0 | 0 | 0 | 0 | 0 | 0 | 0 | 0 | 0 | 0 | 0 | 0 | 0 | 0 | 0 | 0 |
| Oral cavity | | | 0 | 0 | 0 | 0 | 0 | 0 | 0 | 0 | 0 | 0 | 0 | 0 | 0 | 0 | 0 | 0 |
| Skin | | | 0 | 0 | 0 | 0 | 0 | 0 | 0 | 0 | 0 | 0 | 0 | 0 | 0 | 0 | 0 | 0 |
| Subcutaneous tissues | | | 0 | 0 | 0 | 0 | 0 | 0 | 0 | 0 | 0 | 0 | 0 | 0 | 0 | 0 | 0 | 0 |
| Musculoskeletal system | | | 0 | 0 | 0 | 0 | 0 | 0 | 0 | 0 | 0 | 0 | 0 | 0 | 0 | 0 | 0 | 0 |
| Trachea | | 0: No lesion | 0 | 0 | 0 | 0 | 0 | 0 | 0 | 0 | 0 | 0 | 0 | 0 | 0 | 0 | 0 | 0 |
| Lung | 0-1 | 1: Lesion is present | 0 | 0 | 0 | 0 | 0 | 0 | 0 | 0 | 0 | 0 | 0 | 0 | 0 | 0 | 0 | 0 |
| Heart | | | 0 | 1 | 0 | 0 | 0 | 0 | 0 | 0 | 0 | 0 | 0 | 0 | 0 | 0 | 0 | 0 |
| Lymph nodes, tonsil | | | 0 | 0 | 0 | 0 | 0 | 0 | 0 | 0 | 0 | 0 | 0 | 0 | 0 | 0 | 0 | 0 |
| Stomach, intestine | | | 0 | 0 | 0 | 0 | 0 | 0 | 0 | 0 | 0 | 0 | 0 | 0 | 0 | 0 | 0 | 0 |
| Liver | | | 0 | 0 | 0 | 0 | 0 | 0 | 0 | 0 | 0 | 0 | 0 | 0 | 0 | 0 | 0 | 0 |
| Spleen | | | 0 | 0 | 0 | 0 | 0 | 0 | 0 | 0 | 0 | 0 | 0 | 0 | 0 | 0 | 0 | 0 |
| Kidney | | | 0 | 0 | 0 | 0 | 0 | 0 | 0 | 0 | 0 | 0 | 0 | 0 | 0 | 0 | 0 | 0 |
| Brain | | | 0 | 0 | 0 | 0 | 0 | 0 | 0 | 0 | 0 | 0 | 0 | 0 | 0 | 0 | 0 | 0 |
| Body condition | 0-2 | 0: Vertebrae, ribs and bony protuberances only detectable with firm pressure.<br>1: Vertebrae, ribs and bony protuberances prominent.<br>2: Vertebrae, ribs and bony protuberances very prominent | 0 | 0 | 0 | 0 | 0 | 0 | 0 | 0 | 0 | 0 | 0 | 0 | 0 | 0 | 0 | 0 |
| Thoracic cavity | | 0: no lesion | 0 | 1 | 0 | 2 | 0 | 0 | 0 | 0 | 0 | 0 | 0 | 0 | 0 | 0 | 0 | 0 |
| Pericardial cavity | 0-2 | 1: fibrinous serositis in a circumscribed area | 0 | 2 | 0 | 0 | 2 | 0 | 0 | 0 | 2 | 2 | 0 | 0 | 0 | 0 | 0 | 0 |
| Peritoneal cavity | | 2: diffuse/multifocal fibrinous serositis | 0 | 1 | 0 | 0 | 0 | 0 | 0 | 0 | 0 | 0 | 0 | 0 | 0 | 0 | 0 | 0 |
| Stifle | | 0: no lesion | 1 | 1 | 0 | 0 | 0 | 0 | 1 | 1 | 0 | 0 | 0 | 2 | 0 | 0 | 0 | 0 |
| Elbow | | 1: mild serous arthritis with mild joint swelling | 1 | 2 | 0 | 0 | 0 | 0 | 1 | 0 | 1 | 1 | 1 | 0 | 0 | 0 | 0 | 0 |
| Tarsus | 0-2 | 2: severe serous purulent arthritis with pronounced joint swelling | 1 | 2 | 1 | 1 | 0 | 1 | 1 | 0 | 2 | 2 | 0 | 0 | 0 | 0 | 0 | 0 |
| Carpus | | | 2 | 0 | 0 | 0 | 2 | 0 | 0 | 0 | 0 | 2 | 0 | 1 | 0 | 0 | 0 | 0 |
| **Total** | | | **5** | **10** | **1** | **3** | **4** | **1** | **3** | **1** | **5** | **7** | **1** | **3** | **0** | **0** | **0** | **0** |

**Supplementary table 2**

Background information of the strains used for the enzyme-linked immunsorbent assay

| ID | Farm | Country | Tissue | Isolation year |
|---|---|---|---|---|
| MycSu74 | Mohács | Hungary | pericardium | 2016. |
| MycSu78 | Hajdúszoboszló | Hungary | synovial fluid | 2016. |
| MycSu105 | Tiszainoka | Hungary | meninx | 2017. |
| MycSu141 | Bácsalmás | Hungary | serosa | 2019. |
| MycSu199 | Lajoskomárom | Hungary | synovial fluid | 2020. |
| MycSu206 | Tedej | Hungary | pericardium | 2020. |

**Supplementary table 3**

Rectal temperatures measured during the study

| | Group IV-IV | | | | | | Group IV-IP | | | | | | Control Group | | | |
|---|---|---|---|---|---|---|---|---|---|---|---|---|---|---|---|---|
| **Animal ID** | **1** | **2** | **3** | **4** | **5** | **6** | **7** | **8** | **9** | **10** | **11** | **12** | **13** | **14** | **15** | **16** |
| Days | | | | | | | | | | | | | | | | |
| D-2 | 38.3 | 38.6 | 38.8 | 39.1 | 39.6 | 39.2 | 38.1 | 39.7 | 39.4 | 39.5 | 39.0 | 39.0 | 39.6 | 39.2 | 38.5 | 39.6 |
| D-1 | 38.6 | 38.8 | 39.0 | 39.4 | 38.7 | 39.0 | 37.2 | 39.5 | 39.2 | 39.6 | 39.0 | 38.7 | 39.0 | 39.4 | 38.8 | 39.6 |
| D0 | 39.5 | 39.1 | 39.4 | 39.1 | 39.6 | 39.6 | 38.2 | 39.1 | 39.3 | 39.7 | 38.5 | 39.1 | 39.7 | 39.0 | 38.5 | 39.7 |
| D1 | 39.6 | 39.2 | 38.9 | 39.0 | 39.4 | 39.4 | 39.2 | 39.3 | 39.5 | 39.9 | 39.4 | 39.4 | 39.2 | 39.1 | 38.9 | 39.6 |
| D2 | 39.4 | 39.4 | 39.1 | 39.4 | 39.1 | 39.4 | 39.3 | 38.8 | 39.5 | 39.4 | 39.2 | 39.3 | 39.8 | 39.6 | 39.0 | 40.0 |
| D3 | 39.3 | 39.5 | 38.9 | 39.3 | 39.3 | 38.9 | 38.9 | 39.5 | 39.2 | 39.7 | 39.3 | 39.1 | 39.7 | 39.2 | 39.0 | 39.5 |
| D4 | 39.5 | 39.3 | 39.1 | 39.9 | 39.6 | 39.5 | 39.0 | 39.2 | 39.6 | 40.0 | 38.8 | 39.3 | 39.5 | 39.0 | 39.1 | 39.6 |
| D5 | 40.0 | 38.8 | 39.0 | 39.5 | 39.2 | 39.5 | 38.8 | 38.9 | 39.1 | 40.0 | 38.5 | 39.1 | 39.2 | 39.0 | 38.2 | 39.6 |
| D6 | 39.2 | 39.4 | 38.8 | 39.4 | 39.0 | 39.8 | 39.4 | 39.5 | 38.8 | 40.3 | 39.2 | 39.5 | 39.2 | 39.0 | 38.4 | 39.4 |
| D7 | 39.0 | 39.4 | 38.7 | 38.8 | 38.3 | 39.0 | 39.2 | 39.1 | 39.0 | 39.5 | 39.3 | 38.5 | 39.1 | 39.1 | 39.0 | 39.2 |
| D8 | 39.4 | 39.8 | 38.5 | 39.6 | 39.8 | 39.8 | 38.8 | 39.1 | 39.2 | 39.8 | 38.8 | 38.4 | 38.9 | 39.0 | 39.1 | 39.4 |
| D9 | 38.8 | 39.1 | 39.1 | 39.2 | 39.0 | 39.9 | 38.7 | 38.7 | 38.1 | 39.4 | 38.6 | 38.7 | 38.7 | 38.2 | 38.3 | 39.0 |
| D10 | 38.5 | 38.7 | 38.6 | 39.0 | 38.8 | 39.0 | 38.1 | 38.9 | 39.0 | 39.6 | 39.3 | 39.0 | 39.2 | 38.7 | 38.5 | 38.8 |
| D11 | 38.9 | 38.7 | 38.5 | 39.0 | 38.5 | 38.6 | 38.6 | 39.4 | 38.9 | 39.4 | 39.6 | 39.1 | 39.2 | 38.3 | 38.4 | 39.2 |
| D12 | 38.2 | 39.0 | 38.9 | 33.9 | 39.2 | 39.2 | 38.5 | 38.9 | 38.6 | 39.4 | 39.0 | 39.1 | 38.8 | 39.3 | 38.7 | 39.2 |
| D13 | 39.0 | 39.4 | 38.8 | 39.6 | 39.1 | 39.2 | 38.4 | 38.9 | 38.6 | 39.3 | 38.8 | 39.2 | 38.8 | 38.8 | 37.8 | 39.1 |
| D14 | 39.3 | 39.1 | 38.8 | 39.4 | 39.2 | 39.0 | 38.8 | 39.5 | 39.0 | 39.3 | 39.1 | 39.3 | 38.8 | 38.7 | 38.0 | 39.0 |
| D15 | 39.4 | 39.7 | 38.4 | 39.3 | 39.1 | 39.4 | 38.9 | 39.2 | 39.1 | 39.6 | 37.4 | 39.2 | 39.5 | 39.4 | 38.8 | 39.1 |
| D16 | 39.0 | 38.6 | 38.2 | 38.5 | 39.1 | 38.8 | 38.7 | 39.7 | 39.3 | 39.4 | 39.2 | 39.5 | 39.7 | 39.0 | 38.6 | 39.6 |
| D17 | 39.5 | 39.2 | 39.0 | 39.4 | 39.2 | 39.0 | 39.1 | 39.4 | 38.9 | 39.4 | 39.1 | 39.7 | 39.7 | 39.1 | 38.0 | 39.8 |
| D18 | 39.2 | 39.2 | 38.8 | 39.1 | 39.2 | 39.2 | 40.0 | 39.5 | 39.6 | 39.4 | 39.6 | 39.6 | 39.4 | 38.8 | 37.8 | 39.2 |
| D19 | 39.1 | 39.0 | 39.1 | 39.5 | 39.5 | 39.3 | 39.2 | 39.0 | 39.5 | 40.0 | 38.7 | 39.2 | 39.2 | 38.2 | 38.1 | 39.1 |
| D20 | 39.0 | 38.2 | 38.8 | 39.0 | 39.0 | 38.7 | 38.4 | 39.0 | 39.1 | 39.0 | 39.4 | 39.0 | 39.3 | 39.2 | 38.4 | 39.3 |
| D21 | 39.0 | 39.1 | 38.9 | 38.9 | 39.0 | 38.9 | 39.0 | 39.1 | 39.2 | 39.5 | 39.2 | 38.8 | 39.7 | 39.0 | 38.7 | 39.4 |
| D22 | 39.3 | 39.4 | 39.5 | 39.5 | 39.6 | 38.7 | 39.4 | 39.7 | 39.7 | 39.5 | 39.1 | 39.6 | 39.5 | 39.4 | 38.9 | 39.6 |
| D23 | 39.1 | 39.2 | 39.1 | 39.3 | 39.9 | 39.3 | 38.5 | 38.8 | 38.5 | 39.2 | 39.1 | 38.7 | 39.2 | 39.1 | 38.6 | 39.1 |
| D24 | 39.5 | 39.4 | 39.5 | 39.5 | 39.7 | 39.5 | 40.0 | 39.4 | 39.2 | 39.7 | 39.7 | 40.3 | 39.6 | 39.5 | 38.6 | 39.6 |
| D25 | 39.3 | 39.4 | 39.0 | 39.5 | 39.5 | 39.4 | 39.0 | 39.0 | 38.8 | 39.1 | 38.8 | 39.1 | 39.6 | 39.2 | 38.8 | 39.7 |
| D26 | 39.1 | 39.6 | 39.2 | 39.7 | 39.5 | 39.6 | 39.0 | 39.5 | 39.2 | 39.6 | 39.4 | 39.7 | 39.7 | 39.5 | 39.2 | 39.8 |
| D27 | 39.4 | 39.5 | 39.3 | 39.4 | 39.6 | 39.3 | 38.8 | 39.4 | 39.0 | 39.2 | 39.4 | 39.4 | 39.2 | 39.1 | 38.0 | 39.5 |

**Supplementary table 4**

Timescale of apperance of swollen joints

| | Group IV-IV | | | | | | Group IV-IP | | | | | | Control Group | | | |
|---|---|---|---|---|---|---|---|---|---|---|---|---|---|---|---|---|
| **Animal ID** | **1** | **2** | **3** | **4** | **5** | **6** | **7** | **8** | **9** | **10** | **11** | **12** | **13** | **14** | **15** | **16** |
| Days | | | | | | | | | | | | | | | | |
| D6 | | | | | | | | | | + | | | | | | |
| D7 | | | | | | | | | | + | | | | | | |
| D8 | | | + | | | | + | | | + | | | | | | |
| D9 | | | + | | | | + | | + | + | | | | | | |
| D10 | | | + | | | | + | | + | + | | | | | | |
| D11 | + | | + | | | + | + | | + | + | | | | | | |
| D12 | + | | + | | | + | + | | + | + | | | | | | |
| D13 | + | | + | | | + | + | | + | + | | + | | | | |
| D14 | + | | + | + | | + | + | | + | + | + | + | | | | |
| D15 | + | | + | + | | + | + | | + | + | + | + | | | | |
| D16 | + | | + | + | | + | + | + | + | + | + | + | | | | |
| D17 | + | | + | + | | + | + | + | + | + | + | + | | | | |
| D18 | + | | + | + | | + | + | + | + | + | + | + | | | | |
| D19 | + | | + | + | | + | + | + | + | + | + | + | | | | |
| D20 | + | + | + | + | | + | + | + | + | + | + | + | | | | |
| D21 | + | + | + | + | | + | + | + | + | + | + | + | | | | |
| D22 | + | + | + | + | | + | + | + | + | + | + | + | | | | |
| D23 | + | + | + | + | | + | + | + | + | + | + | + | | | | |
| D24 | + | + | + | + | | + | + | + | + | + | + | + | | | | |
| D25 | + | + | + | + | | + | + | + | + | + | + | + | | | | |
| D26 | + | + | + | + | | + | + | + | + | + | + | + | | | | |
| D27 | + | + | + | + | | + | + | + | + | + | + | + | | | | |

+: at least one swollen joint

**Supplementary table 5**

Results of weight measurement in kilograms

| | Group IV-IV | | | | | | Group IV-IP | | | | | | Control Group | | | |
|---|---|---|---|---|---|---|---|---|---|---|---|---|---|---|---|---|
| **Animal ID** | **1** | **2** | **3** | **4** | **5** | **6** | **7** | **8** | **9** | **10** | **11** | **12** | **13** | **14** | **15** | **16** |
| Weeks | | | | | | | | | | | | | | | | |
| W0 | 11.0 | 11.0 | 8.5 | 12.0 | 10.0 | 11.0 | 9.0 | 12.0 | 11.0 | 10.0 | 11.0 | 9.0 | 12.0 | 9.5 | 10.0 | 10.0 |
| W1 | 13.0 | 10.0 | 10.0 | 14.0 | 11.0 | 12.0 | 8.0 | 13.0 | 10.0 | 10.0 | 11.0 | 9.0 | 11.0 | 9.0 | 10.5 | 15.0 |
| W1.5 | 11.0 | 10.0 | 9.5 | 15.0 | 10.0 | 13.0 | 11.0 | 13.0 | 11.0 | 12.0 | 13.0 | 10.0 | 14.0 | 10.0 | 11.0 | 13.0 |
| W2 | 14.0 | 13.0 | 11.0 | 15.0 | 12.0 | 11.0 | 12.0 | 17.0 | 13.0 | 13.0 | 10.0 | 13.0 | 13.0 | 10.0 | 12.0 | 13.0 |
| W2.5 | 14.0 | 10.0 | 11.0 | 16.0 | 11.0 | 14.0 | 11.0 | 15.0 | 12.0 | 10.5 | 14.0 | 11.5 | 15.5 | 10.0 | 10.0 | 14.0 |
| W3 | 16.0 | 14.0 | 14.0 | 19.0 | 14.0 | 16.0 | 15.0 | 18.0 | 15.0 | 14.0 | 16.0 | 12.0 | 18.0 | 12.0 | 16.0 | 18.0 |
| W3.5 | 16.0 | 14.0 | 14.0 | 19.0 | 13.0 | 18.0 | 14.0 | 19.0 | 15.0 | 14.0 | 16.0 | 13.0 | 20.0 | 14.0 | 15.0 | 19.0 |
| W4 | 17.0 | 17.0 | 17.0 | 22.0 | 14.0 | 19.0 | 14.0 | 19.0 | 14.0 | 14.0 | 16.0 | 13.0 | 23.0 | 14.0 | 16.0 | 22.0 |
| W4.5 | 19.0 | 15.0 | 15.0 | 24.0 | 15.0 | 20.0 | 15.0 | 20.0 | 15.0 | 15.0 | 17.0 | 14.0 | 26.0 | 18.0 | 19.0 | 25.0 |

**Supplementary table 6**

*Mycoplasma hyorhinis* specific PCR and isolation results of the nasal swabs

| | Group IV-IV | | | | | | Group IV-IP | | | | | | Control Group | | | |
|---|---|---|---|---|---|---|---|---|---|---|---|---|---|---|---|---|
| **Animal ID** | **1** | **2** | **3** | **4** | **5** | **6** | **7** | **8** | **9** | **10** | **11** | **12** | **13** | **14** | **15** | **16** |
| **Sampling time** | | | | | | | | | | | | | | | | |
| D-2 | - | - | - | - | - | - | - | - | - | - | - | - | - | - | - | - |
| D5 | - | - | - | - | - | - | - | - | + | ++ | - | - | - | - | - | - |
| D8 | - | + | - | - | - | - | - | - | + | + | - | - | - | - | - | - |
| D12 | - | - | - | (+) | - | - | - | - | - | + | - | - | - | - | - | - |
| D15 | - | - | - | + | + | - | - | - | - | - | - | - | - | - | - | - |
| D19 | - | - | - | - | - | - | - | - | - | - | - | - | - | - | - | - |
| D21 | - | - | - | + | - | - | - | - | - | - | - | - | - | - | - | - |
| D26 | - | + | - | + | - | - | - | - | - | - | - | - | - | - | - | - |

(+): positive Mycoplasma isolation with negative PCR

+: PCR positivity without positive Mycoplasma isolation

++: PCR positivity and positive Mycoplasma isolation

**Supplementary table 7**

*Mycoplasma hyorhinis* specific PCR and isolation results of the necropsy swab samples

| | Group IV-IV | | | | | | Group IV-IP | | | | | | Control Group | | | |
|---|---|---|---|---|---|---|---|---|---|---|---|---|---|---|---|---|
| **Animal ID** | **1** | **2** | **3** | **4** | **5** | **6** | **7** | **8** | **9** | **10** | **11** | **12** | **13** | **14** | **15** | **16** |
| Pericardium | - | + | - | - | - | - | - | - | - | - | - | - | - | - | - | - |
| Pleura | - | - | - | + | - | - | - | - | - | - | - | - | - | - | - | - |
| Peritoneum | - | + | - | - | - | - | - | - | - | - | - | - | - | - | - | - |
| Stifle | + | + | - | - | - | - | + | ++ | - | - | - | - | - | - | - | - |
| Elbow | + | + | - | + | - | ++ | + | ++ | + | - | + | - | - | - | - | - |
| Tarsus | + | + | + | ++ | - | + | + | ++ | + | + | - | - | - | - | - | - |
| Carpus | + | + | + | - | + | - | - | - | - | ++ | - | - | - | - | - | - |
| Conjunctiva | - | - | - | - | - | - | - | - | - | - | - | - | - | - | - | - |
| Meninx | - | - | - | - | - | - | - | - | - | - | - | - | - | - | - | - |
| Lung | - | + | - | - | - | - | - | - | - | - | - | - | - | - | - | - |

+: PCR positivity without positive Mycoplasma isolation

++: PCR positivity and positive Mycoplasma isolation

**Supplementary table 8**

Source of isolation and repeat numbers of the re-isolates

| | | | Repeat numbers | | | | | |
|---|---|---|---|---|---|---|---|---|
| **Group** | **Source** | **Animal** | **Mhr205** | **Mhr396** | **Mhr438** | **Mhr441** | **Mhr442** | **Mhr444** |
| MycSu160 masterseed | | | 1 | 2 | 0 | 15 | 0 | 46 |
| Group IV-IV | nasal cavity | A | 1 | 2 | 0 | 15 | 0 | 46 |
| Group IV-IV | tarsus | A | 1 | 2 | 0 | 15 | 0 | 46 |
| Group IV-IV | elbow | B | 1 | 2 | 0 | 15 | 0 | 46 |
| Group IV-IP | carpus | A | 1 | 2 | 0 | 15 | 0 | 42 |
| Group IV-IP | nasal cavity | A | 1 | 2 | 0 | 15 | 0 | 46 |
| Group IV-IP | elbow | B | 1 | 2 | 0 | 15 | 0 | 46 |
| Group IV-IP | tarsus | B | 1 | 2 | 0 | 15 | 0 | 48 |
| Group IV-IP | stifle | B | 1 | 2 | 0 | 15 | 0 | 46 |

**Supplementary table 9**

Histopathological scores of the study groups

| | Group IV-IV | | | | | | Group IV-IP | | | | | | Control Group | | | |
|---|---|---|---|---|---|---|---|---|---|---|---|---|---|---|---|---|
| **Animal ID** | **1** | **2** | **3** | **4** | **5** | **6** | **7** | **8** | **9** | **10** | **11** | **12** | **13** | **14** | **15** | **16** |
| Pericardium | 0 | 3 | 0 | 0 | 3 | 1 | 0 | 0 | 3 | 3 | 0 | 0 | 0 | 0 | 0 | 0 |
| Pleura | 0 | 2 | 0 | 3 | 0 | 2 | 2 | 1 | 2 | 1 | 0 | 0 | 0 | 0 | 0 | 0 |
| Peritoneum | 0 | 2 | 0 | 0 | 0 | 0 | 0 | 0 | 1 | 0 | 0 | 0 | 0 | 0 | 0 | 0 |
| Stifle | 0 | 3 | 0 | 0 | 0 | 0 | 3 | 3 | 3 | 3 | 0 | 0 | 0 | 0 | 0 | 0 |
| Elbow | 0 | 3 | 0 | 0 | 0 | 2 | 1 | 0 | 3 | 3 | 0 | 0 | 0 | 0 | 0 | 0 |
| Tarsus | 2 | 3 | 2 | 3 | 0 | 2 | 1 | 0 | 3 | 3 | 0 | 0 | 0 | 0 | 0 | 0 |
| Carpus | 3 | 3 | 0 | 0 | 3 | 0 | 0 | 0 | 1 | 3 | 0 | 0 | 0 | 0 | 0 | 0 |
| Conjunctiva | 0 | 0 | 2 | 0 | 0 | 0 | 0 | 0 | 1 | 0 | 0 | 0 | 0 | 0 | 0 | 0 |
| Brain | 0 | 0 | 0 | 0 | 0 | 0 | 0 | 0 | 0 | 0 | 0 | 0 | 0 | 0 | 0 | 0 |
| Lung | 0 | 0 | 0 | 0 | 0 | 0 | 0 | 0 | 0 | 0 | 0 | 0 | 0 | 0 | 0 | 0 |
| Nasal mucosal membrane | 0 | 1 | 0 | 0 | 0 | 0 | 0 | 0 | 0 | 0 | 0 | 0 | 0 | 0 | 0 | 0 |
| **Total** | **5** | **20** | **4** | **6** | **6** | **7** | **7** | **4** | **17** | **16** | **0** | **0** | **0** | **0** | **0** | **0** |

0: no lesion

1: mild lesion

2: moderate lesion

3: severe lesion

**Supplementary table 10**

Anti-*Mycoplasma hyorhinis* enzyme-linked immunsorbent assay results given in S/P%

| | Group IV-IV | | | | | | Group IV-IP | | | | | | Control Group | | | |
|---|---|---|---|---|---|---|---|---|---|---|---|---|---|---|---|---|
| **Animal ID** | **1** | **2** | **3** | **4** | **5** | **6** | **7** | **8** | **9** | **10** | **11** | **12** | **13** | **14** | **15** | **16** |
| **Sampling time** | | | | | | | | | | | | | | | | |
| D-2 | 14.29 | 11.16 | 11.80 | 14.58 | 10.98 | 17.16 | 16.79 | 18.27 | 12.08 | 12.18 | 14.30 | 14.02 | 13.28 | 9.23 | 15.40 | 12.64 |
| D5 | 46.49 | 34.13 | 20.11 | 33.03 | 31.65 | 37.45 | 37.64 | 39.61 | 43.01 | 62.08 | 28.23 | 25.00 | 21.13 | 12.55 | 17.71 | 16.14 |
| D8 | 63.37 | 33.79 | 30.16 | 31.55 | 35.61 | 29.36 | 25.74 | 35.06 | 36.16 | 63.84 | 32.47 | 29.98 | 20.30 | 12.82 | 25.83 | 20.39 |
| D12 | 54.94 | 68.93 | 38.22 | 32.79 | 37.10 | 35.40 | 36.11 | 12.69 | 56.69 | 50.92 | 15.30 | 15.16 | 5.29 | 1.76 | 3.95 | 2.19 |
| D15 | 70.80 | 44.36 | 33.78 | 47.53 | 44.15 | 28.84 | 89.77 | 23.91 | 75.18 | 52.61 | 36.46 | 32.37 | 9.52 | 4.16 | 6.49 | 2.68 |
| D19 | 83.57 | 45.98 | 32.44 | 60.86 | 54.51 | 28.28 | 111.36 | 35.15 | 97.04 | 72.78 | 34.27 | 26.23 | 10.01 | 2.75 | 9.59 | 2.54 |
| D21 | 65.94 | 52.47 | 37.02 | 46.33 | 47.95 | 58.74 | 81.10 | 43.30 | 85.19 | 61.14 | 29.19 | 31.45 | 10.72 | 4.87 | 5.50 | 0.28 |
| D25 | 122.71 | 76.73 | 53.10 | 65.79 | 51.48 | 79.48 | 99.37 | 68.27 | 101.20 | 107.83 | 49.22 | 39.77 | 17.14 | 6.14 | 12.98 | 1.69 |
| D28 | 133.00 | 84.34 | 68.40 | 71.79 | 69.96 | 86.53 | 92.17 | 74.33 | 115.80 | 118.12 | 59.10 | 47.04 | 18.41 | 7.69 | 10.44 | 2.82 |

S/P% > 40 are highlighted green and considered positive
