## Supplementary figures and images for "Establishment of a *Mycoplasma hyorhinis* challenge model in five-week-old piglets"

### Supplementary figure 1

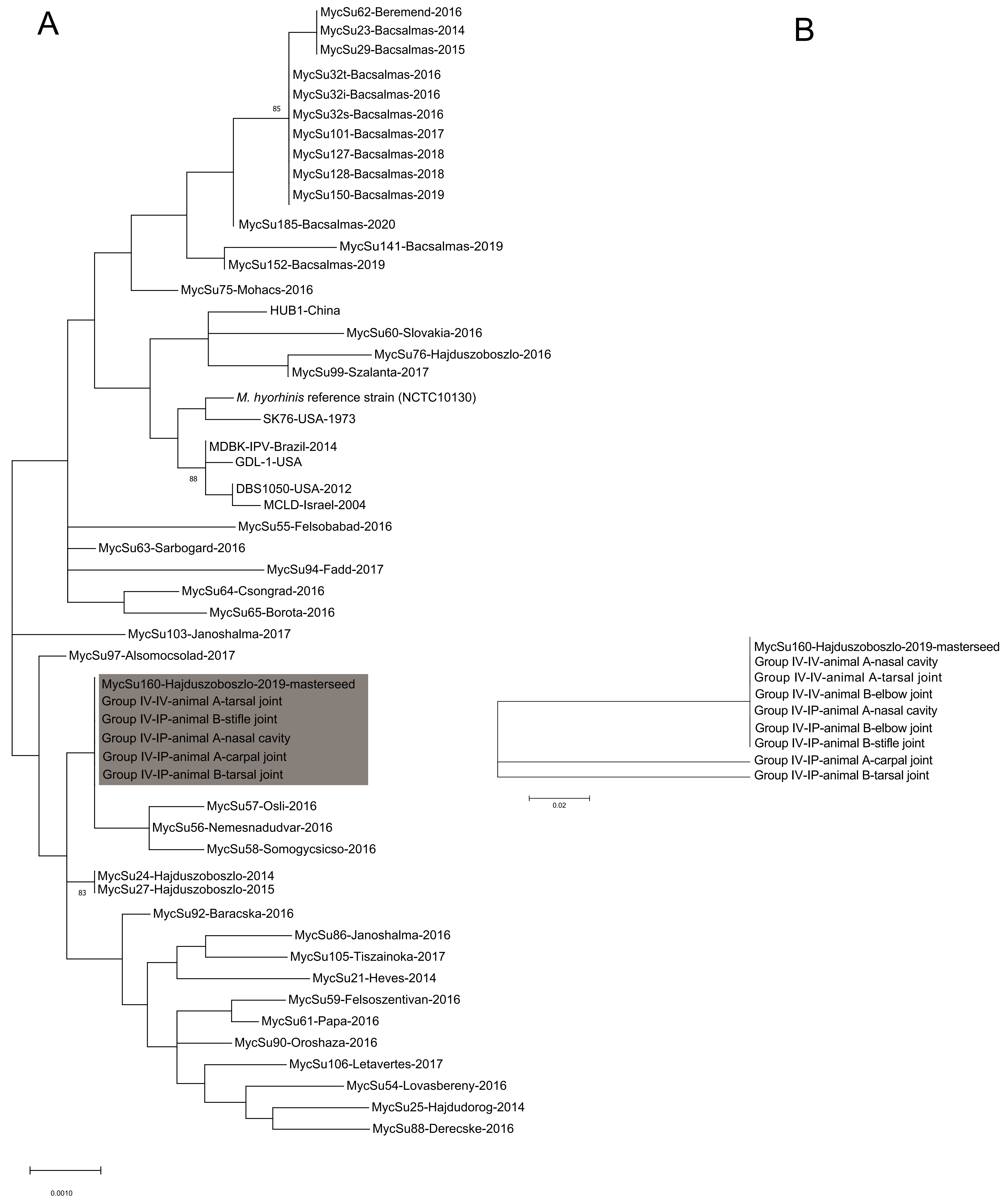
