## Supplementary data 1 for "Establishment of a *Mycoplasma hyorhinis* challenge model in five-week-old piglets"

**a) Statistical analysis of daily weight gain**

| | Number of animals | Mean | SD |
|---|---|---|---|
| Group IV-IV | 6 | 223 | 86.90 |
| Group IV-IP | 6 | 170 | 41.00 |
| Control | 4 | 350 | 99.00 |

Shapiro-Wilk normality test

Null hypothesis: the distribution of the data is not significantly different form normal distribution.

W=0.89, p-value=0.06

one-way ANOVA

| | DF | Sum Sq | Mean Sq | F-value | Pr (>F) |
|---|---|---|---|---|---|
| Group | 2 | 79067 | 39533 | 6.80 | **<0.01** |
| Residuals | 13 | 75533 | 5810 | | |

Tukey multiple comparisons of means

| Comparison | diff | lwr | upr | adjusted p-value |
|---|---|---|---|---|
| Group IV-IP-Group IV-IV | -53.33 | -169.54 | -62.87 | 0.47 |
| Control-Group IV-IV | 126.67 | -3.25 | 256.58 | **0.05** |
| Control-Group IV-IP | 180.00 | 50.08 | 309.92 | **<0.01** |

**b) Statistical analysis of gross pathological scores of joint lesions**

| | Number of animals | Median | IQR |
|---|---|---|---|
| Group IV-IV | 6 | 1.50 | 3.25 |
| Group IV-IP | 6 | 3.00 | 1.50 |
| Control | 4 | 0.00 | 0.00 |

Kruskal-Wallis test

| | |
|---|---|
| chi-squared | 9.03 |
| DF | 2 |
| p-value | **0.01** |

Dunn-test

| Comparison | Z-value | unadjusted p-value | adjusted p-value |
|---|---|---|---|
| Group IV-IV-Group IV-IP | -0.25 | 0.80 | 1.00 |
| Group IV-IV- Control | 2.57 | 0.01 | **0.03** |
| Group IV-IP- Control | 2.79 | 0.01 | **0.02** |

**c) Statistical analysis of gross pathological scores of serosa of pericardium, pleura and peritoneum**

| | Number of animals | Median | IQR |
|---|---|---|---|
| Group IV-IV | 6 | 0.50 | 1.75 |
| Group IV-IP | 6 | 0.00 | 1.50 |
| Control | 4 | 0.00 | 0.00 |

| Kruskal-Wallis test | |
|---|---|
| chi-squared | 2.54 |
| DF | 2 |
| p-value | 0.28 |

**d) Statistical analysis of total gross pathological scores**

| | Number of animals | Median | IQR |
|---|---|---|---|
| Group IV-IV | 6 | 3.00 | 3.50 |
| Group IV-IP | 6 | 3.00 | 3.00 |
| Control | 4 | 0.00 | 0.00 |

| Kruskal-Wallis test | |
|---|---|
| chi-squared | 8.76 |
| DF | 2 |
| p-value | **0.01** |

Dunn-test

| Comparison | Z-value | unadjusted p-value | adjusted p-value |
|---|---|---|---|
| Group IV-IV-Group IV-IP | 0.06 | 0.95 | 1.00 |
| Group IV-IV- Control | 2.67 | 0.01 | **0.02** |
| Group IV-IP- Control | 2.62 | 0.01 | **0.03** |

**e) Statistical analysis of joint histology scores**

| | Number of animals | Median | IQR |
|---|---|---|---|
| Group IV-IV | 6 | 3.50 | 1.75 |
| Group IV-IP | 6 | 3.50 | 7.75 |
| Control | 4 | 0.00 | 0.00 |

| Kruskal-Wallis test | |
|---|---|
| chi-squared | 6.51 |
| DF | 2 |
| p-value | **0.04** |

| Dunn-test | | | |
|---|---|---|---|
| Comparison | Z-value | unadjusted p-value | adjusted p-value |
| Group IV-IV-Group IV-IP | 0.50 | 0.62 | 1.00 |
| Group IV-IV- Control | 2.46 | 0.01 | **0.04** |
| Group IV-IP- Control | 2.01 | 0.04 | 0.13 |

**f) Statistical analysis of the histology scores of serosa of pericardium, pleura and peritoneum**

| | Number of animals | Median | IQR |
|---|---|---|---|
| Group IV-IV | 6 | 3.00 | 0.75 |
| Group IV-IP | 6 | 2.00 | 3.50 |
| Control | 4 | 0.00 | 0.00 |

| Kruskal-Wallis test | |
|---|---|
| chi-squared | 5.48 |
| DF | 2 |
| p-value | 0.06 |

**g) Statistical analysis of total histology scores**

| | Number of animals | Median | IQR |
|---|---|---|---|
| Group IV-IV | 6 | 6.00 | 1.50 |
| Group IV-IP | 6 | 5.50 | 12.80 |
| Control | 4 | 0.00 | 0.00 |

| Kruskal-Wallis test | |
|---|---|
| chi-squared | 6.62 |
| DF | 2 |
| p-value | **0.04** |

| Dunn-test | | | |
|---|---|---|---|
| Comparison | Z-value | unadjusted p-value | adjusted p-value |
| Group IV-IV-Group IV-IP | 0.62 | 0.53 | 1.00 |
| Group IV-IV- Control | 2.51 | 0.01 | **0.04** |
| Group IV-IP- Control | 1.95 | 0.51 | 0.15 |

**h) Statistical analysis of ELISA S/P% from the last sampling point**

| | Number of animals | Mean | SD |
|---|---|---|---|
| Group IV-IV | 6 | 85.70 | 24.40 |
| Group IV-IP | 6 | 84.40 | 29.40 |
| Control | 4 | 9.84 | 6.52 |

Shapiro-Wilk normality test

Null hypothesis: the distribution of the data is not significantly different form normal distribution.

W=0.94, p-value=0.38

one-way ANOVA

| | DF | Sum Sq | Mean Sq | F-value | Pr (>F) |
|---|---|---|---|---|---|
| Group | 2 | 16974 | 8487 | 14.85 | **<0.01** |
| Residuals | 13 | 7428 | 571 | | |

Tukey multiple comparisons of means

| Comparison | diff | lwr | upr | adjusted p-value |
|---|---|---|---|---|
| Group IV-IP-Group IV-IV | -1.25 | -37.69 | 35.20 | 0.99 |
| Control-Group IV-IV | -75.83 | -116.57 | -35.09 | **<0.01** |
| Control-Group IV-IP | -74.59 | -115.33 | -33.84 | **<0.01** |
