## Supplementary data 2 for "Establishment of a *Mycoplasma hyorhinis* challenge model in five-week-old piglets"

```
              ....|....| ....|....| ....|....| ....|....| ....|....| ....|....|
                  5         15         25         35         45         55
Group_IV-I    ATAGAAAAAT TCGTACAAGA TATGCACCTA GTCCAACTGG ATATTTACAC ATAGGTGGAG
Group_IV-I    ATAGAAAAAT TCGTACAAGA TATGCACCTA GTCCAACTGG ATATTTACAC ATAGGTGGAG
Group_IV-I    ATAGAAAAAT TCGTACAAGA TATGCACCTA GTCCAACTGG ATATTTACAC ATAGGTGGAG
Group_IV-I    ATAGAAAAAT TCGTACAAGA TATGCACCTA GTCCAACTGG ATATTTACAC ATAGGTGGAG
Group_IV-I    ATAGAAAAAT TCGTACAAGA TATGCACCTA GTCCAACTGG ATATTTACAC ATAGGTGGAG
MycSu160_m    ATAGAAAAAT TCGTACAAGA TATGCACCTA GTCCAACTGG ATATTTACAC ATAGGTGGAG

              ....|....| ....|....| ....|....| ....|....| ....|....| ....|....|
                  65         75         85         95        105        115
Group_IV-I    CAAGAACTGC GTTATTTAAT TATTTATTTG CCAAACATTT TTTAGGTGAT TTCGTATTAA
Group_IV-I    CAAGAACTGC GTTATTTAAT TATTTATTTG CCAAACATTT TTTAGGTGAT TTCGTATTAA
Group_IV-I    CAAGAACTGC GTTATTTAAT TATTTATTTG CCAAACATTT TTTAGGTGAT TTCGTATTAA
Group_IV-I    CAAGAACTGC GTTATTTAAT TATTTATTTG CCAAACATTT TTTAGGTGAT TTCGTATTAA
Group_IV-I    CAAGAACTGC GTTATTTAAT TATTTATTTG CCAAACATTT TTTAGGTGAT TTCGTATTAA
MycSu160_m    CAAGAACTGC GTTATTTAAT TATTTATTTG CCAAACATTT TTTAGGTGAT TTCGTATTAA

              ....|....| ....|....| ....|....| ....|....| ....|....| ....|....|
                 125        135        145        155        165        175
Group_IV-I    GAATTGAAGA TACAGATGTT GCAAGAAATG TAGTAGGTGG AGAAAAATCT CAATTAGATA
Group_IV-I    GAATTGAAGA TACAGATGTT GCAAGAAATG TAGTAGGTGG AGAAAAATCT CAATTAGATA
Group_IV-I    GAATTGAAGA TACAGATGTT GCAAGAAATG TAGTAGGTGG AGAAAAATCT CAATTAGATA
Group_IV-I    GAATTGAAGA TACAGATGTT GCAAGAAATG TAGTAGGTGG AGAAAAATCT CAATTAGATA
Group_IV-I    GAATTGAAGA TACAGATGTT GCAAGAAATG TAGTAGGTGG AGAAAAATCT CAATTAGATA
MycSu160_m    GAATTGAAGA TACAGATGTT GCAAGAAATG TAGTAGGTGG AGAAAAATCT CAATTAGATA

              ....|....| ....|....| ....|....| ....|....| ....|....| ....|....|
                 185        195        205        215        225        235
Group_IV-I    ATTTATTGTG GTTGGGCATT ATTCCAGATG AATACCCAAC TAAAGAAAAT TCAAAATATG
Group_IV-I    ATTTATTGTG GTTGGGCATT ATTCCAGATG AATACCCAAC TAAAGAAAAT TCAAAATATG
Group_IV-I    ATTTATTGTG GTTGGGCATT ATTCCAGATG AATACCCAAC TAAAGAAAAT TCAAAATATG
Group_IV-I    ATTTATTGTG GTTGGGCATT ATTCCAGATG AATACCCAAC TAAAGAAAAT TCAAAATATG
Group_IV-I    ATTTATTGTG GTTGGGCATT ATTCCAGATG AATACCCAAC TAAAGAAAAT TCAAAATATG
MycSu160_m    ATTTATTGTG GTTGGGCATT ATTCCAGATG AATACCCAAC TAAAGAAAAT TCAAAATATG

              ....|....| ....|....| ....|....| ....|....| ....|....| ....|....|
                 245        255        265        275        285        295
Group_IV-I    GTTTTTATCG TCAATCACAA AAATTAAAAA GATATTGAGA TTTAGCACAT CAACTAATAG
Group_IV-I    GTTTTTATCG TCAATCACAA AAATTAAAAA GATATTGAGA TTTAGCACAT CAACTAATAG
Group_IV-I    GTTTTTATCG TCAATCACAA AAATTAAAAA GATATTGAGA TTTAGCACAT CAACTAATAG
Group_IV-I    GTTTTTATCG TCAATCACAA AAATTAAAAA GATATTGAGA TTTAGCACAT CAACTAATAG
Group_IV-I    GTTTTTATCG TCAATCACAA AAATTAAAAA GATATTGAGA TTTAGCACAT CAACTAATAG
MycSu160_m    GTTTTTATCG TCAATCACAA AAATTAAAAA GATATTGAGA TTTAGCACAT CAACTAATAG

              ....|....| ....|....| ....|....| ....|....| ....|....| ....|....|
                 305        315        325        335        345        355
Group_IV-I    CACAAAAAAA AGCTTATTTA GCATTTGATT CACCAGAAGA AATTCAACTA CAAAAACAAG
Group_IV-I    CACAAAAAAA AGCTTATTTA GCATTTGATT CACCAGAAGA AATTCAACTA CAAAAACAAG
```

```
Group_IV-I   CACAAAAAAA AGCTTATTTA GCATTTGATT CACCAGAAGA AATTCAACTA CAAAAACAAG
Group_IV-I   CACAAAAAAA AGCTTATTTA GCATTTGATT CACCAGAAGA AATTCAACTA CAAAAACAAG
Group_IV-I   CACAAAAAAA AGCTTATTTA GCATTTGATT CACCAGAAGA AATTCAACTA CAAAAACAAG
MycSu160_m   CACAAAAAAA AGCTTATTTA GCATTTGATT CACCAGAAGA AATTCAACTA CAAAAACAAG

             ....|....| ....|....| ....|....| ....|....| ....|....| ....|....|
                 365        375        385        395        405        415
Group_IV-I   AACAAGAAAA AAAAGGAATT TTTAGTTTTA GATACGATAG AAACTGGTTA ATTGGTATTC
Group_IV-I   AACAAGAAAA AAAAGGAATT TTTAGTTTTA GATACGATAG AAACTGGTTA ATTGGTATTC
Group_IV-I   AACAAGAAAA AAAAGGAATT TTTAGTTTTA GATACGATAG AAACTGGTTA ATTGGTATTC
Group_IV-I   AACAAGAAAA AAAAGGAATT TTTAGTTTTA GATACGATAG AAACTGGTTA ATTGGTATTC
Group_IV-I   AACAAGAAAA AAAAGGAATT TTTAGTTTTA GATACGATAG AAACTGGTTA ATTGGTATTC
MycSu160_m   AACAAGAAAA AAAAGGAATT TTTAGTTTTA GATACGATAG AAACTGGTTA ATTGGTATTC

             ....|....| ....|....| ....|....| ....|....| ....|....| ....|....|
                 425        435        445        455        465        475
Group_IV-I   CAACAGATGA TGCGATTTTA ATTTCTGCTA AAAATGGTAC AAATATTGAA GCTGTTTTAG
Group_IV-I   CAACAGATGA TGCGATTTTA ATTTCTGCTA AAAATGGTAC AAATATTGAA GCTGTTTTAG
Group_IV-I   CAACAGATGA TGCGATTTTA ATTTCTGCTA AAAATGGTAC AAATATTGAA GCTGTTTTAG
Group_IV-I   CAACAGATGA TGCGATTTTA ATTTCTGCTA AAAATGGTAC AAATATTGAA GCTGTTTTAG
Group_IV-I   CAACAGATGA TGCGATTTTA ATTTCTGCTA AAAATGGTAC AAATATTGAA GCTGTTTTAG
MycSu160_m   CAACAGATGA TGCGATTTTA ATTTCTGCTA AAAATGGTAC AAATATTGAA GCTGTTTTAG

             ....|....| ....|....| ....|....| ....|....| ....|....| ....|....|
                 485        495        505        515        525        535
Group_IV-I   AAGCAATTAT TAATAAAATT CCAGCTCCAA AAGTGGATGA AAATGAAAAA GAATTAAAAG
Group_IV-I   AAGCAATTAT TAATAAAATT CCAGCTCCAA AAGTGGATGA AAATGAAAAA GAATTAAAAG
Group_IV-I   AAGCAATTAT TAATAAAATT CCAGCTCCAA AAGTGGATGA AAATGAAAAA GAATTAAAAG
Group_IV-I   AAGCAATTAT TAATAAAATT CCAGCTCCAA AAGTGGATGA AAATGAAAAA GAATTAAAAG
Group_IV-I   AAGCAATTAT TAATAAAATT CCAGCTCCAA AAGTGGATGA AAATGAAAAA GAATTAAAAG
MycSu160_m   AAGCAATTAT TAATAAAATT CCAGCTCCAA AAGTGGATGA AAATGAAAAA GAATTAAAAG

             ....|....| ....|....| ....|....| ....|....| ....|....| ....|....|
                 545        555        565        575        585        595
Group_IV-I   CTCTAGTTTT TGATTCTTAT TTTGATATTT ACAGAGGAGT AATTATTTTA GTAAGAATTG
Group_IV-I   CTCTAGTTTT TGATTCTTAT TTTGATATTT ACAGAGGAGT AATTATTTTA GTAAGAATTG
Group_IV-I   CTCTAGTTTT TGATTCTTAT TTTGATATTT ACAGAGGAGT AATTATTTTA GTAAGAATTG
Group_IV-I   CTCTAGTTTT TGATTCTTAT TTTGATATTT ACAGAGGAGT AATTATTTTA GTAAGAATTG
Group_IV-I   CTCTAGTTTT TGATTCTTAT TTTGATATTT ACAGAGGAGT AATTATTTTA GTAAGAATTG
MycSu160_m   CTCTAGTTTT TGATTCTTAT TTTGATATTT ACAGAGGAGT AATTATTTTA GTAAGAATTG

             ....|....| ....|....| ....|....| ....|....| ....|....| ....|....|
                 605        615        625        635        645        655
Group_IV-I   TTTCAGGTTC TATTAAAGTG GGTGACGAAT TTAAGTTCAT GGCAAATGGT AAGAAGTTTG
Group_IV-I   TTTCAGGTTC TATTAAAGTG GGTGACGAAT TTAAGTTCAT GGCAAATGGT AAGAAGTTTG
Group_IV-I   TTTCAGGTTC TATTAAAGTG GGTGACGAAT TTAAGTTCAT GGCAAATGGT AAGAAGTTTG
Group_IV-I   TTTCAGGTTC TATTAAAGTG GGTGACGAAT TTAAGTTCAT GGCAAATGGT AAGAAGTTTG
Group_IV-I   TTTCAGGTTC TATTAAAGTG GGTGACGAAT TTAAGTTCAT GGCAAATGGT AAGAAGTTTG
MycSu160_m   TTTCAGGTTC TATTAAAGTG GGTGACGAAT TTAAGTTCAT GGCAAATGGT AAGAAGTTTG

             ....|....| ....|....| ....|....| ....|....| ....|....| ....|....|
```

```
                  665        675        685        695        705        715
Group_IV-I   GTGTGATTGA ATTAGGGGTA AGAACTCCAA AAGAGCTTAA AAAAGAAGAG CTTGTTGCAG
Group_IV-I   GTGTGATTGA ATTAGGGGTA AGAACTCCAA AAGAGCTTAA AAAAGAAGAG CTTGTTGCAG
Group_IV-I   GTGTGATTGA ATTAGGGGTA AGAACTCCAA AAGAGCTTAA AAAAGAAGAG CTTGTTGCAG
Group_IV-I   GTGTGATTGA ATTAGGGGTA AGAACTCCAA AAGAGCTTAA AAAAGAAGAG CTTGTTGCAG
Group_IV-I   GTGTGATTGA ATTAGGGGTA AGAACTCCAA AAGAGCTTAA AAAAGAAGAG CTTGTTGCAG
MycSu160_m   GTGTGATTGA ATTAGGGGTA AGAACTCCAA AAGAGCTTAA AAAAGAAGAG CTTGTTGCAG

             ....|....| ....|....| ....|....| ....|....| ....|....| ....|....|
                  725        735        745        755        765        775
Group_IV-I   GTGAAGTGGG TTGAATTGCA GCTTCAATTA GAAACGCAAA GGATGTTTCT GTTGGTGATA
Group_IV-I   GTGAAGTGGG TTGAATTGCA GCTTCAATTA GAAACGCAAA GGATGTTTCT GTTGGTGATA
Group_IV-I   GTGAAGTGGG TTGAATTGCA GCTTCAATTA GAAACGCAAA GGATGTTTCT GTTGGTGATA
Group_IV-I   GTGAAGTGGG TTGAATTGCA GCTTCAATTA GAAACGCAAA GGATGTTTCT GTTGGTGATA
Group_IV-I   GTGAAGTGGG TTGAATTGCA GCTTCAATTA GAAACGCAAA GGATGTTTCT GTTGGTGATA
MycSu160_m   GTGAAGTGGG TTGAATTGCA GCTTCAATTA GAAACGCAAA GGATGTTTCT GTTGGTGATA

             ....|....| ....|....| ....|....| ....|....| ....|....| ....|....|
                  785        795        805        815        825        835
Group_IV-I   CTATTACTTT AGTTGCAAAT CCAGCAAAAG TAGCACTTTC TGGTTATAAA AAATTAAAGC
Group_IV-I   CTATTACTTT AGTTGCAAAT CCAGCAAAAG TAGCACTTTC TGGTTATAAA AAATTAAAGC
Group_IV-I   CTATTACTTT AGTTGCAAAT CCAGCAAAAG TAGCACTTTC TGGTTATAAA AAATTAAAGC
Group_IV-I   CTATTACTTT AGTTGCAAAT CCAGCAAAAG TAGCACTTTC TGGTTATAAA AAATTAAAGC
Group_IV-I   CTATTACTTT AGTTGCAAAT CCAGCAAAAG TAGCACTTTC TGGTTATAAA AAATTAAAGC
MycSu160_m   CTATTACTTT AGTTGCAAAT CCAGCAAAAG TAGCACTTTC TGGTTATAAA AAATTAAAGC

             ....|....| ....|....| ....|....| ....|....| ....|....| ....|....|
                  845        855        865        875        885        895
Group_IV-I   CCGTAGTTTA TACAAGAGAA ATTCCAAATG CTTCACAAAG AAGTAAAAGA CACTTAGATG
Group_IV-I   CCGTAGTTTA TACAAGAGAA ATTCCAAATG CTTCACAAAG AAGTAAAAGA CACTTAGATG
Group_IV-I   CCGTAGTTTA TACAAGAGAA ATTCCAAATG CTTCACAAAG AAGTAAAAGA CACTTAGATG
Group_IV-I   CCGTAGTTTA TACAAGAGAA ATTCCAAATG CTTCACAAAG AAGTAAAAGA CACTTAGATG
Group_IV-I   CCGTAGTTTA TACAAGAGAA ATTCCAAATG CTTCACAAAG AAGTAAAAGA CACTTAGATG
MycSu160_m   CCGTAGTTTA TACAAGAGAA ATTCCAAATG CTTCACAAAG AAGTAAAAGA CACTTAGATG

             ....|....| ....|....| ....|....| ....|....| ....|....| ....|....|
                  905        915        925        935        945        955
Group_IV-I   AAAACGGAAT TGTCCGTATT GGATCAGAAG TTAGTGCAGG TGATATTTTA GTAGGTAGAA
Group_IV-I   AAAACGGAAT TGTCCGTATT GGATCAGAAG TTAGTGCAGG TGATATTTTA GTAGGTAGAA
Group_IV-I   AAAACGGAAT TGTCCGTATT GGATCAGAAG TTAGTGCAGG TGATATTTTA GTAGGTAGAA
Group_IV-I   AAAACGGAAT TGTCCGTATT GGATCAGAAG TTAGTGCAGG TGATATTTTA GTAGGTAGAA
Group_IV-I   AAAACGGAAT TGTCCGTATT GGATCAGAAG TTAGTGCAGG TGATATTTTA GTAGGTAGAA
MycSu160_m   AAAACGGAAT TGTCCGTATT GGATCAGAAG TTAGTGCAGG TGATATTTTA GTAGGTAGAA

             ....|....| ....|....| ....|....| ....|....| ....|....| ....|....|
                  965        975        985        995       1005       1015
Group_IV-I   CTTCACCAAA AGGAGAAGAT AATCCAACAC CTGAAGAAAA ATTAATGAAC GCAATTTGAG
Group_IV-I   CTTCACCAAA AGGAGAAGAT AATCCAACAC CTGAAGAAAA ATTAATGAAC GCAATTTGAG
Group_IV-I   CTTCACCAAA AGGAGAAGAT AATCCAACAC CTGAAGAAAA ATTAATGAAC GCAATTTGAG
Group_IV-I   CTTCACCAAA AGGAGAAGAT AATCCAACAC CTGAAGAAAA ATTAATGAAC GCAATTTGAG
Group_IV-I   CTTCACCAAA AGGAGAAGAT AATCCAACAC CTGAAGAAAA ATTAATGAAC GCAATTTGAG
```

```
MycSu160_m   CTTCACCAAA AGGAGAAGAT AATCCAACAC CTGAAGAAAA ATTAATGAAC GCAATTTGAG

             ....|....| ....|....| ....|....| ....|....| ....|....| ....|....|
                 1025       1035       1045       1055       1065       1075
Group_IV-I   GTAAAAAAGC CTCATCTCAG AAGGATACTT CATTAAAAGT TAAACACGGT GATGGTGGTA
Group_IV-I   GTAAAAAAGC CTCATCTCAG AAGGATACTT CATTAAAAGT TAAACACGGT GATGGTGGTA
Group_IV-I   GTAAAAAAGC CTCATCTCAG AAGGATACTT CATTAAAAGT TAAACACGGT GATGGTGGTA
Group_IV-I   GTAAAAAAGC CTCATCTCAG AAGGATACTT CATTAAAAGT TAAACACGGT GATGGTGGTA
Group_IV-I   GTAAAAAAGC CTCATCTCAG AAGGATACTT CATTAAAAGT TAAACACGGT GATGGTGGTA
MycSu160_m   GTAAAAAAGC CTCATCTCAG AAGGATACTT CATTAAAAGT TAAACACGGT GATGGTGGTA

             ....|....| ....|....| ....|....| ....|....| ....|....| ....|....|
                 1085       1095       1105       1115       1125       1135
Group_IV-I   CAGTTATTGA TGTGCAAATT CTTTCAAGAA CTCGTGAAGA TAATTTAGAA GACGGTGTTG
Group_IV-I   CAGTTATTGA TGTGCAAATT CTTTCAAGAA CTCGTGAAGA TAATTTAGAA GACGGTGTTG
Group_IV-I   CAGTTATTGA TGTGCAAATT CTTTCAAGAA CTCGTGAAGA TAATTTAGAA GACGGTGTTG
Group_IV-I   CAGTTATTGA TGTGCAAATT CTTTCAAGAA CTCGTGAAGA TAATTTAGAA GACGGTGTTG
Group_IV-I   CAGTTATTGA TGTGCAAATT CTTTCAAGAA CTCGTGAAGA TAATTTAGAA GACGGTGTTG
MycSu160_m   CAGTTATTGA TGTGCAAATT CTTTCAAGAA CTCGTGAAGA TAATTTAGAA GACGGTGTTG

             ....|....| ....|....| ....|....| ....|....| ....|....| ....|....|
                 1145       1155       1165       1175       1185       1195
Group_IV-I   AAAAAATCAT CAAAGTCTTT ATCGCTCAAA AAAGAAAAAT TAAAGTTGGG GATAAAATGG
Group_IV-I   AAAAAATCAT CAAAGTCTTT ATCGCTCAAA AAAGAAAAAT TAAAGTTGGG GATAAAATGG
Group_IV-I   AAAAAATCAT CAAAGTCTTT ATCGCTCAAA AAAGAAAAAT TAAAGTTGGG GATAAAATGG
Group_IV-I   AAAAAATCAT CAAAGTCTTT ATCGCTCAAA AAAGAAAAAT TAAAGTTGGG GATAAAATGG
Group_IV-I   AAAAAATCAT CAAAGTCTTT ATCGCTCAAA AAAGAAAAAT TAAAGTTGGG GATAAAATGG
MycSu160_m   AAAAAATCAT CAAAGTCTTT ATCGCTCAAA AAAGAAAAAT TAAAGTTGGG GATAAAATGG

             ....|....| ....|....| ....|....| ....|....| ....|....| ....|....|
                 1205       1215       1225       1235       1245       1255
Group_IV-I   CCGGAAGACA CGGAAACAAA GGAGTTATTT CCTTAGTTTT ACCAGTTGAA GATATGCCAT
Group_IV-I   CCGGAAGACA CGGAAACAAA GGAGTTATTT CCTTAGTTTT ACCAGTTGAA GATATGCCAT
Group_IV-I   CCGGAAGACA CGGAAACAAA GGAGTTATTT CCTTAGTTTT ACCAGTTGAA GATATGCCAT
Group_IV-I   CCGGAAGACA CGGAAACAAA GGAGTTATTT CCTTAGTTTT ACCAGTTGAA GATATGCCAT
Group_IV-I   CCGGAAGACA CGGAAACAAA GGAGTTATTT CCTTAGTTTT ACCAGTTGAA GATATGCCAT
MycSu160_m   CCGGAAGACA CGGAAACAAA GGAGTTATTT CCTTAGTTTT ACCAGTTGAA GATATGCCAT

             ....|....| ....|....| ....|....| ....|....| ....|....| ....|....|
                 1265       1275       1285       1295       1305       1315
Group_IV-I   TTTTAGAAGA TGGAACACCA ATTGATATTC TTTTAAATCC ACAAGGTATT CCTTCACGTA
Group_IV-I   TTTTAGAAGA TGGAACACCA ATTGATATTC TTTTAAATCC ACAAGGTATT CCTTCACGTA
Group_IV-I   TTTTAGAAGA TGGAACACCA ATTGATATTC TTTTAAATCC ACAAGGTATT CCTTCACGTA
Group_IV-I   TTTTAGAAGA TGGAACACCA ATTGATATTC TTTTAAATCC ACAAGGTATT CCTTCACGTA
Group_IV-I   TTTTAGAAGA TGGAACACCA ATTGATATTC TTTTAAATCC ACAAGGTATT CCTTCACGTA
MycSu160_m   TTTTAGAAGA TGGAACACCA ATTGATATTC TTTTAAATCC ACAAGGTATT CCTTCACGTA

             ....|....| ....|....| ....|....| ....|....| ....|....| ....|....|
                 1325       1335       1345       1355       1365       1375
Group_IV-I   TGAACTGAAA GATATATTCT TTCAATTAAC AAATGAACAG CTGTAAAAAA TGAAATCCAA
Group_IV-I   TGAACTGAAA GATATATTCT TTCAATTAAC AAATGAACAG CTGTAAAAAA TGAAATCCAA
```

```
Group_IV-I   TGAACTGAAA GATATATTCT TTCAATTAAC AAATGAACAG CTGTAAAAAA TGAAATCCAA
Group_IV-I   TGAACTGAAA GATATATTCT TTCAATTAAC AAATGAACAG CTGTAAAAAA TGAAATCCAA
Group_IV-I   TGAACTGAAA GATATATTCT TTCAATTAAC AAATGAACAG CTGTAAAAAA TGAAATCCAA
MycSu160_m   TGAACTGAAA GATATATTCT TTCAATTAAC AAATGAACAG CTGTAAAAAA TGAAATCCAA

             ....|....| ....|....| ....|....| ....|....| ....|....| ....|....|
                 1385       1395       1405       1415       1425       1435
Group_IV-I   AAAAATCTAA ACGAAACAGT TGAACTTGAT CCAAAAAATT CACTAGTTAT TATGATGAAA
Group_IV-I   AAAAATCTAA ACGAAACAGT TGAACTTGAT CCAAAAAATT CACTAGTTAT TATGATGAAA
Group_IV-I   AAAAATCTAA ACGAAACAGT TGAACTTGAT CCAAAAAATT CACTAGTTAT TATGATGAAA
Group_IV-I   AAAAATCTAA ACGAAACAGT TGAACTTGAT CCAAAAAATT CACTAGTTAT TATGATGAAA
Group_IV-I   AAAAATCTAA ACGAAACAGT TGAACTTGAT CCAAAAAATT CACTAGTTAT TATGATGAAA
MycSu160_m   AAAAATCTAA ACGAAACAGT TGAACTTGAT CCAAAAAATT CACTAGTTAT TATGATGAAA

             ....|....| ....|....| ....|....| ....|....| ....|....| ....|....|
                 1445       1455       1465       1475       1485       1495
Group_IV-I   TCAGGAGCTA GATCAAATAT GTCTAACTTC GTTCAATTAG CTGGAATGCG TGGATTGATG
Group_IV-I   TCAGGAGCTA GATCAAATAT GTCTAACTTC GTTCAATTAG CTGGAATGCG TGGATTGATG
Group_IV-I   TCAGGAGCTA GATCAAATAT GTCTAACTTC GTTCAATTAG CTGGAATGCG TGGATTGATG
Group_IV-I   TCAGGAGCTA GATCAAATAT GTCTAACTTC GTTCAATTAG CTGGAATGCG TGGATTGATG
Group_IV-I   TCAGGAGCTA GATCAAATAT GTCTAACTTC GTTCAATTAG CTGGAATGCG TGGATTGATG
MycSu160_m   TCAGGAGCTA GATCAAATAT GTCTAACTTC GTTCAATTAG CTGGAATGCG TGGATTGATG

             ....|....| ....|....| ....|....| ....|....| ....|....| ....|....|
                 1505       1515       1525       1535       1545       1555
Group_IV-I   GCAAATAACG TTAAAGCTCT AAAGGTAGAT GCTGAAAATG AAAGAGTTGT TCGTTCAATT
Group_IV-I   GCAAATAACG TTAAAGCTCT AAAGGTAGAT GCTGAAAATG AAAGAGTTGT TCGTTCAATT
Group_IV-I   GCAAATAACG TTAAAGCTCT AAAGGTAGAT GCTGAAAATG AAAGAGTTGT TCGTTCAATT
Group_IV-I   GCAAATAACG TTAAAGCTCT AAAGGTAGAT GCTGAAAATG AAAGAGTTGT TCGTTCAATT
Group_IV-I   GCAAATAACG TTAAAGCTCT AAAGGTAGAT GCTGAAAATG AAAGAGTTGT TCGTTCAATT
MycSu160_m   GCAAATAACG TTAAAGCTCT AAAGGTAGAT GCTGAAAATG AAAGAGTTGT TCGTTCAATT

             ....|....| ....|....| ....|....| ....|....| ....|....| ....|....|
                 1565       1575       1585       1595       1605       1615
Group_IV-I   GTTGAAGTGC CAGTTAAATC TTCATTCTTA GAAGGTCTTA CATCATTTGA GTTCTATTCT
Group_IV-I   GTTGAAGTGC CAGTTAAATC TTCATTCTTA GAAGGTCTTA CATCATTTGA GTTCTATTCT
Group_IV-I   GTTGAAGTGC CAGTTAAATC TTCATTCTTA GAAGGTCTTA CATCATTTGA GTTCTATTCT
Group_IV-I   GTTGAAGTGC CAGTTAAATC TTCATTCTTA GAAGGTCTTA CATCATTTGA GTTCTATTCT
Group_IV-I   GTTGAAGTGC CAGTTAAATC TTCATTCTTA GAAGGTCTTA CATCATTTGA GTTCTATTCT
MycSu160_m   GTTGAAGTGC CAGTTAAATC TTCATTCTTA GAAGGTCTTA CATCATTTGA GTTCTATTCT

             ....|....| ....|....| ....|....| ....|....| ....|....| ....|....|
                 1625       1635       1645       1655       1665       1675
Group_IV-I   TCAACACACG GTGCTCGAAA AGGGCTTACC GATACAGCGC TTAACACAGC TAAATCAGGT
Group_IV-I   TCAACACACG GTGCTCGAAA AGGGCTTACC GATACAGCGC TTAACACAGC TAAATCAGGT
Group_IV-I   TCAACACACG GTGCTCGAAA AGGGCTTACC GATACAGCGC TTAACACAGC TAAATCAGGT
Group_IV-I   TCAACACACG GTGCTCGAAA AGGGCTTACC GATACAGCGC TTAACACAGC TAAATCAGGT
Group_IV-I   TCAACACACG GTGCTCGAAA AGGGCTTACC GATACAGCGC TTAACACAGC TAAATCAGGT
MycSu160_m   TCAACACACG GTGCTCGAAA AGGGCTTACC GATACAGCGC TTAACACAGC TAAATCAGGT

             ....|....| ....|....| ....|....| ....|....| ....|....| ....|....|
```

```
                 1685       1695       1705       1715       1725       1735
Group_IV-I   TATTTAACTC GTCGTTTAGT TGATGTAGCA CAAAACATTG TTGTTACTCA AAATGATTGT
Group_IV-I   TATTTAACTC GTCGTTTAGT TGATGTAGCA CAAAACATTG TTGTTACTCA AAATGATTGT
Group_IV-I   TATTTAACTC GTCGTTTAGT TGATGTAGCA CAAAACATTG TTGTTACTCA AAATGATTGT
Group_IV-I   TATTTAACTC GTCGTTTAGT TGATGTAGCA CAAAACATTG TTGTTACTCA AAATGATTGT
Group_IV-I   TATTTAACTC GTCGTTTAGT TGATGTAGCA CAAAACATTG TTGTTACTCA AAATGATTGT
MycSu160_m   TATTTAACTC GTCGTTTAGT TGATGTAGCA CAAAACATTG TTGTTACTCA AAATGATTGT

             ....|....| ....|....| ....|....| ....|....| ....|....| ....|....|
                 1745       1755       1765       1775       1785       1795
Group_IV-I   TTTTCTGACT TTGGATTTGT AGTAAAACAA ATTATTGATA CTAAAACAAA CACAACAATT
Group_IV-I   TTTTCTGACT TTGGATTTGT AGTAAAACAA ATTATTGATA CTAAAACAAA CACAACAATT
Group_IV-I   TTTTCTGACT TTGGATTTGT AGTAAAACAA ATTATTGATA CTAAAACAAA CACAACAATT
Group_IV-I   TTTTCTGACT TTGGATTTGT AGTAAAACAA ATTATTGATA CTAAAACAAA CACAACAATT
Group_IV-I   TTTTCTGACT TTGGATTTGT AGTAAAACAA ATTATTGATA CTAAAACAAA CACAACAATT
MycSu160_m   TTTTCTGACT TTGGATTTGT AGTAAAACAA ATTATTGATA CTAAAACAAA CACAACAATT

             ....|....| ....|....| ....|....| ....|....| ....|....| ....|....|
                 1805       1815       1825       1835       1845       1855
Group_IV-I   GTTCCTTTAG CAGAAAGAAT TGAAGGTAGA TTTTTAAACA AAGATGTAGT AAGTCCGGAT
Group_IV-I   GTTCCTTTAG CAGAAAGAAT TGAAGGTAGA TTTTTAAACA AAGATGTAGT AAGTCCGGAT
Group_IV-I   GTTCCTTTAG CAGAAAGAAT TGAAGGTAGA TTTTTAAACA AAGATGTAGT AAGTCCGGAT
Group_IV-I   GTTCCTTTAG CAGAAAGAAT TGAAGGTAGA TTTTTAAACA AAGATGTAGT AAGTCCGGAT
Group_IV-I   GTTCCTTTAG CAGAAAGAAT TGAAGGTAGA TTTTTAAACA AAGATGTAGT AAGTCCGGAT
MycSu160_m   GTTCCTTTAG CAGAAAGAAT TGAAGGTAGA TTTTTAAACA AAGATGTAGT AAGTCCGGAT

             ....|....| ....|....| ....|....| ....|....| ....|....| ....|....|
                 1865       1875       1885       1895       1905       1915
Group_IV-I   GGTGAAGTAC TAGCACAAAC AGGTGATTTT TTAAAAATAA AGATTCTATT ACTTCTCAGT
Group_IV-I   GGTGAAGTAC TAGCACAAAC AGGTGATTTT TTAAAAATAA AGATTCTATT ACTTCTCAGT
Group_IV-I   GGTGAAGTAC TAGCACAAAC AGGTGATTTT TTAAAAATAA AGATTCTATT ACTTCTCAGT
Group_IV-I   GGTGAAGTAC TAGCACAAAC AGGTGATTTT TTAAAAATAA AGATTCTATT ACTTCTCAGT
Group_IV-I   GGTGAAGTAC TAGCACAAAC AGGTGATTTT TTAAAAATAA AGATTCTATT ACTTCTCAGT
MycSu160_m   GGTGAAGTAC TAGCACAAAC AGGTGATTTT TTAAAAATAA AGATTCTATT ACTTCTCAGT

             ....|....| ....|....| ....|....| ....|....| ....|....| ....|....|
                 1925       1935       1945       1955       1965       1975
Group_IV-I   ATCTTTTAGG CATTAAAAAA ATTGAAATTC CTTCAAAAAG AAGAATGGGT AATGGGCAAT
Group_IV-I   ATCTTTTAGG CATTAAAAAA ATTGAAATTC CTTCAAAAAG AAGAATGGGT AATGGGCAAT
Group_IV-I   ATCTTTTAGG CATTAAAAAA ATTGAAATTC CTTCAAAAAG AAGAATGGGT AATGGGCAAT
Group_IV-I   ATCTTTTAGG CATTAAAAAA ATTGAAATTC CTTCAAAAAG AAGAATGGGT AATGGGCAAT
Group_IV-I   ATCTTTTAGG CATTAAAAAA ATTGAAATTC CTTCAAAAAG AAGAATGGGT AATGGGCAAT
MycSu160_m   ATCTTTTAGG CATTAAAAAA ATTGAAATTC CTTCAAAAAG AAGAATGGGT AATGGGCAAT

             ....|....| ....|....| ....|....| ....|....| ....|....| ....|....|
                 1985       1995       2005       2015       2025       2035
Group_IV-I   TTATAGTAAT TGAAGGTGCT AAAGAAAACA ATTTAAAAAA CTTAAAAGTA GAAATACCAT
Group_IV-I   TTATAGTAAT TGAAGGTGCT AAAGAAAACA ATTTAAAAAA CTTAAAAGTA GAAATACCAT
Group_IV-I   TTATAGTAAT TGAAGGTGCT AAAGAAAACA ATTTAAAAAA CTTAAAAGTA GAAATACCAT
Group_IV-I   TTATAGTAAT TGAAGGTGCT AAAGAAAACA ATTTAAAAAA CTTAAAAGTA GAAATACCAT
Group_IV-I   TTATAGTAAT TGAAGGTGCT AAAGAAAACA ATTTAAAAAA CTTAAAAGTA GAAATACCAT
```

```
MycSu160_m   TTATAGTAAT TGAAGGTGCT AAAGAAAACA ATTTAAAAAA CTTAAAAGTA GAAATACCAT

             ....|....| ....|....| ....|....| ....|....| ....|....| ....|....|
                 2045       2055       2065       2075       2085       2095
Group_IV-I   TGGGTAAATT TGTAGCTGTA ACTGGAGTTT CTGGTTCAGG AAAATCTACT TTAGTAAATG
Group_IV-I   TGGGTAAATT TGTAGCTGTA ACTGGAGTTT CTGGTTCAGG AAAATCTACT TTAGTAAATG
Group_IV-I   TGGGTAAATT TGTAGCTGTA ACTGGAGTTT CTGGTTCAGG AAAATCTACT TTAGTAAATG
Group_IV-I   TGGGTAAATT TGTAGCTGTA ACTGGAGTTT CTGGTTCAGG AAAATCTACT TTAGTAAATG
Group_IV-I   TGGGTAAATT TGTAGCTGTA ACTGGAGTTT CTGGTTCAGG AAAATCTACT TTAGTAAATG
MycSu160_m   TGGGTAAATT TGTAGCTGTA ACTGGAGTTT CTGGTTCAGG AAAATCTACT TTAGTAAATG

             ....|....| ....|....| ....|....| ....|....| ....|....| ....|....|
                 2105       2115       2125       2135       2145       2155
Group_IV-I   AAATTTTAGT TAATGGAATT GTAAAACATC TAACAAATCC TTCACAAAAG GTAGGAAAAC
Group_IV-I   AAATTTTAGT TAATGGAATT GTAAAACATC TAACAAATCC TTCACAAAAG GTAGGAAAAC
Group_IV-I   AAATTTTAGT TAATGGAATT GTAAAACATC TAACAAATCC TTCACAAAAG GTAGGAAAAC
Group_IV-I   AAATTTTAGT TAATGGAATT GTAAAACATC TAACAAATCC TTCACAAAAG GTAGGAAAAC
Group_IV-I   AAATTTTAGT TAATGGAATT GTAAAACATC TAACAAATCC TTCACAAAAG GTAGGAAAAC
MycSu160_m   AAATTTTAGT TAATGGAATT GTAAAACATC TAACAAATCC TTCACAAAAG GTAGGAAAAC

             ....|....| ....|....| ....|....| ....|....| ....|....| ....|....|
                 2165       2175       2185       2195       2205       2215
Group_IV-I   ATTCGCAAAT AAAAGGAATG TTTAATCTTG ATAAAATAGT TTCCATTTCA CAATCACCAA
Group_IV-I   ATTCGCAAAT AAAAGGAATG TTTAATCTTG ATAAAATAGT TTCCATTTCA CAATCACCAA
Group_IV-I   ATTCGCAAAT AAAAGGAATG TTTAATCTTG ATAAAATAGT TTCCATTTCA CAATCACCAA
Group_IV-I   ATTCGCAAAT AAAAGGAATG TTTAATCTTG ATAAAATAGT TTCCATTTCA CAATCACCAA
Group_IV-I   ATTCGCAAAT AAAAGGAATG TTTAATCTTG ATAAAATAGT TTCCATTTCA CAATCACCAA
MycSu160_m   ATTCGCAAAT AAAAGGAATG TTTAATCTTG ATAAAATAGT TTCCATTTCA CAATCACCAA

             ....|....| ....|....| ....|....| ....|....| ....|....| ....|....|
                 2225       2235       2245       2255       2265       2275
Group_IV-I   TTGGAAGAAC GCCTCGTTCT AATCCAGCAA CTTATACTTC GGTTTTTAAC GATATTCGAG
Group_IV-I   TTGGAAGAAC GCCTCGTTCT AATCCAGCAA CTTATACTTC GGTTTTTAAC GATATTCGAG
Group_IV-I   TTGGAAGAAC GCCTCGTTCT AATCCAGCAA CTTATACTTC GGTTTTTAAC GATATTCGAG
Group_IV-I   TTGGAAGAAC GCCTCGTTCT AATCCAGCAA CTTATACTTC GGTTTTTAAC GATATTCGAG
Group_IV-I   TTGGAAGAAC GCCTCGTTCT AATCCAGCAA CTTATACTTC GGTTTTTAAC GATATTCGAG
MycSu160_m   TTGGAAGAAC GCCTCGTTCT AATCCAGCAA CTTATACTTC GGTTTTTAAC GATATTCGAG

             ....|....| ....|....| ....|....| ....|....| ....|....| ....|....|
                 2285       2295       2305       2315       2325       2335
Group_IV-I   ATATTTTTGC TTCAGTTGAG TTAGCTCGAG CTAGAGGTTA TCAAAAAGGT CATTTTTCTT
Group_IV-I   ATATTTTTGC TTCAGTTGAG TTAGCTCGAG CTAGAGGTTA TCAAAAAGGT CATTTTTCTT
Group_IV-I   ATATTTTTGC TTCAGTTGAG TTAGCTCGAG CTAGAGGTTA TCAAAAAGGT CATTTTTCTT
Group_IV-I   ATATTTTTGC TTCAGTTGAG TTAGCTCGAG CTAGAGGTTA TCAAAAAGGT CATTTTTCTT
Group_IV-I   ATATTTTTGC TTCAGTTGAG TTAGCTCGAG CTAGAGGTTA TCAAAAAGGT CATTTTTCTT
MycSu160_m   ATATTTTTGC TTCAGTTGAG TTAGCTCGAG CTAGAGGTTA TCAAAAAGGT CATTTTTCTT

             ....|....| ....|....| ....|....| ....|....| ....|....| ....|....|
                 2345       2355       2365       2375       2385       2395
Group_IV-I   TTAATTTAGC AATTGGACGT TGTGATAAAT GTCAAGGTGA TGGTTCAATT AAAATTGAAA
Group_IV-I   TTAATTTAGC AATTGGACGT TGTGATAAAT GTCAAGGTGA TGGTTCAATT AAAATTGAAA
```

```
Group_IV-I   TTAATTTAGC AATTGGACGT TGTGATAAAT GTCAAGGTGA TGGTTCAATT AAAATTGAAA
Group_IV-I   TTAATTTAGC AATTGGACGT TGTGATAAAT GTCAAGGTGA TGGTTCAATT AAAATTGAAA
Group_IV-I   TTAATTTAGC AATTGGACGT TGTGATAAAT GTCAAGGTGA TGGTTCAATT AAAATTGAAA
MycSu160_m   TTAATTTAGC AATTGGACGT TGTGATAAAT GTCAAGGTGA TGGTTCAATT AAAATTGAAA

             ....|....| ....|....| ....|....| ....|....| ....|....| ....|....|
                 2405       2415       2425       2435       2445       2455
Group_IV-I   TGCATTTTGT TTTAAACAAA ACAATTTCCA AATATATGAA TAAATATAAT TTTTCATTAA
Group_IV-I   TGCATTTTGT TTTAAACAAA ACAATTTCCA AATATATGAA TAAATATAAT TTTTCATTAA
Group_IV-I   TGCATTTTGT TTTAAACAAA ACAATTTCCA AATATATGAA TAAATATAAT TTTTCATTAA
Group_IV-I   TGCATTTTGT TTTAAACAAA ACAATTTCCA AATATATGAA TAAATATAAT TTTTCATTAA
Group_IV-I   TGCATTTTGT TTTAAACAAA ACAATTTCCA AATATATGAA TAAATATAAT TTTTCATTAA
MycSu160_m   TGCATTTTGT TTTAAACAAA ACAATTTCCA AATATATGAA TAAATATAAT TTTTCATTAA

             ....|....| ....|....| ....|....| ....|....| ....|....| ....|....|
                 2465       2475       2485       2495       2505       2515
Group_IV-I   TTTCTAAATC TATAAACGAG TTTGTTTTTG AAGATTTTTC ATCTTGATAT ATTGAATTAA
Group_IV-I   TTTCTAAATC TATAAACGAG TTTGTTTTTG AAGATTTTTC ATCTTGATAT ATTGAATTAA
Group_IV-I   TTTCTAAATC TATAAACGAG TTTGTTTTTG AAGATTTTTC ATCTTGATAT ATTGAATTAA
Group_IV-I   TTTCTAAATC TATAAACGAG TTTGTTTTTG AAGATTTTTC ATCTTGATAT ATTGAATTAA
Group_IV-I   TTTCTAAATC TATAAACGAG TTTGTTTTTG AAGATTTTTC ATCTTGATAT ATTGAATTAA
MycSu160_m   TTTCTAAATC TATAAACGAG TTTGTTTTTG AAGATTTTTC ATCTTGATAT ATTGAATTAA

             ....|....| ....|....| ....|....| ....|....| ....|....| ....|....|
                 2525       2535       2545       2555       2565       2575
Group_IV-I   ATAAATTACA TCAAAATGGA TATCATTTAA GAAAGGTTCT TAAAAAAGTT TTAATTGTTC
Group_IV-I   ATAAATTACA TCAAAATGGA TATCATTTAA GAAAGGTTCT TAAAAAAGTT TTAATTGTTC
Group_IV-I   ATAAATTACA TCAAAATGGA TATCATTTAA GAAAGGTTCT TAAAAAAGTT TTAATTGTTC
Group_IV-I   ATAAATTACA TCAAAATGGA TATCATTTAA GAAAGGTTCT TAAAAAAGTT TTAATTGTTC
Group_IV-I   ATAAATTACA TCAAAATGGA TATCATTTAA GAAAGGTTCT TAAAAAAGTT TTAATTGTTC
MycSu160_m   ATAAATTACA TCAAAATGGA TATCATTTAA GAAAGGTTCT TAAAAAAGTT TTAATTGTTC

             ....|....| ....|....| ....|....| ....|....| ....|....| ....|....|
                 2585       2595       2605       2615       2625       2635
Group_IV-I   TACATCCTTT TATTCCATTT TTAACTGATT ATTTATTTAA AGAAATCTTT AACGAAGAGC
Group_IV-I   TACATCCTTT TATTCCATTT TTAACTGATT ATTTATTTAA AGAAATCTTT AACGAAGAGC
Group_IV-I   TACATCCTTT TATTCCATTT TTAACTGATT ATTTATTTAA AGAAATCTTT AACGAAGAGC
Group_IV-I   TACATCCTTT TATTCCATTT TTAACTGATT ATTTATTTAA AGAAATCTTT AACGAAGAGC
Group_IV-I   TACATCCTTT TATTCCATTT TTAACTGATT ATTTATTTAA AGAAATCTTT AACGAAGAGC
MycSu160_m   TACATCCTTT TATTCCATTT TTAACTGATT ATTTATTTAA AGAAATCTTT AACGAAGAGC

             ....|....| ....|....| ....|....| ....|....| ....|....| ....|....|
                 2645       2655       2665       2675       2685       2695
Group_IV-I   TACTTGAGCA AAAAAGATTA ATGTTTAGAA ATTATAAAAA CACAGAAAAA ATTGATAAAG
Group_IV-I   TACTTGAGCA AAAAAGATTA ATGTTTAGAA ATTATAAAAA CACAGAAAAA ATTGATAAAG
Group_IV-I   TACTTGAGCA AAAAAGATTA ATGTTTAGAA ATTATAAAAA CACAGAAAAA ATTGATAAAG
Group_IV-I   TACTTGAGCA AAAAAGATTA ATGTTTAGAA ATTATAAAAA CACAGAAAAA ATTGATAAAG
Group_IV-I   TACTTGAGCA AAAAAGATTA ATGTTTAGAA ATTATAAAAA CACAGAAAAA ATTGATAAAG
MycSu160_m   TACTTGAGCA AAAAAGATTA ATGTTTAGAA ATTATAAAAA CACAGAAAAA ATTGATAAAG

             ....|....| ....|....| ....|....| ....|....| ....|....| ....|....|
```

```
                 2705      2715      2725      2735      2745      2755
Group_IV-I   TTATTGAAAT TGTTTCTGAA CTTAGAAAAT ATCGTGAAAA ACATAATATT TCTAAAAAAG
Group_IV-I   TTATTGAAAT TGTTTCTGAA CTTAGAAAAT ATCGTGAAAA ACATAATATT TCTAAAAAAG
Group_IV-I   TTATTGAAAT TGTTTCTGAA CTTAGAAAAT ATCGTGAAAA ACATAATATT TCTAAAAAAG
Group_IV-I   TTATTGAAAT TGTTTCTGAA CTTAGAAAAT ATCGTGAAAA ACATAATATT TCTAAAAAAG
Group_IV-I   TTATTGAAAT TGTTTCTGAA CTTAGAAAAT ATCGTGAAAA ACATAATATT TCTAAAAAAG
MycSu160_m   TTATTGAAAT TGTTTCTGAA CTTAGAAAAT ATCGTGAAAA ACATAATATT TCTAAAAAAG

             ....|....| ....|....| ....|....| ....|....| ....|....| ....|....|
                 2765      2775      2785      2795      2805      2815
Group_IV-I   AAAAATTACA ATATTGAATT AAAAATAATT CTTTAGATTC TGATTCATTG AACTTAATCA
Group_IV-I   AAAAATTACA ATATTGAATT AAAAATAATT CTTTAGATTC TGATTCATTG AACTTAATCA
Group_IV-I   AAAAATTACA ATATTGAATT AAAAATAATT CTTTAGATTC TGATTCATTG AACTTAATCA
Group_IV-I   AAAAATTACA ATATTGAATT AAAAATAATT CTTTAGATTC TGATTCATTG AACTTAATCA
Group_IV-I   AAAAATTACA ATATTGAATT AAAAATAATT CTTTAGATTC TGATTCATTG AACTTAATCA
MycSu160_m   AAAAATTACA ATATTGAATT AAAAATAATT CTTTAGATTC TGATTCATTG AACTTAATCA

             ....|....| ....|....| ....|....| ....|....| ....|....| ....|....|
                 2825      2835      2845      2855      2865      2875
Group_IV-I   ATAAATTAGC AATTGCTGAA ATCTTTGAAA ATAATGTTTC TATTTTAAAA ACCAATAATT
Group_IV-I   ATAAATTAGC AATTGCTGAA ATCTTTGAAA ATAATGTTTC TATTTTAAAA ACCAATAATT
Group_IV-I   ATAAATTAGC AATTGCTGAA ATCTTTGAAA ATAATGTTTC TATTTTAAAA ACCAATAATT
Group_IV-I   ATAAATTAGC AATTGCTGAA ATCTTTGAAA ATAATGTTTC TATTTTAAAA ACCAATAATT
Group_IV-I   ATAAATTAGC AATTGCTGAA ATCTTTGAAA ATAATGTTTC TATTTTAAAA ACCAATAATT
MycSu160_m   ATAAATTAGC AATTGCTGAA ATCTTTGAAA ATAATGTTTC TATTTTAAAA ACCAATAATT

             ....|....| ....|....| ....|....| ....|....| ....|....| ....|....|
                 2885      2895      2905      2915      2925      2935
Group_IV-I   TCGAAATTTT CTTAAATCTT TCAGAATCCA AGCAAAATGA AGAACAATTA AGAATAGAAA
Group_IV-I   TCGAAATTTT CTTAAATCAT TCAGAATCCA AGCAAAATGA AGAACAATTA AGAATAGAAA
Group_IV-I   TCGAAATTTT CTTAAATCTT TCAGAATCCA AGCAAAATGA AGAACAATTA AGAATAGAAA
Group_IV-I   TCGAAATTTT CTTAAATCTT TCAGAATCCA AGCAAAATGA AGAACAATTA AGAATAGAAA
Group_IV-I   TCGAAATTTT CTTAAATCTT TCAGAATCCA AGCAAAATGA AGAACAATTA AGAATAGAAA
MycSu160_m   TCGAAATTTT CTTAAATCTT TCAGAATCCA AGCAAAATGA AGAACAATTA AGAATAGAAA

             ....|....| ....|....| ....|....| ....|....| ....|....| ....|....|
                 2945      2955      2965      2975      2985      2995
Group_IV-I   AAGAAATTAA ATATTTAGAA TCCGAAGTCC AAAGATCTTC TGCAATCTTG TCAAATCCTA
Group_IV-I   AAGAAATTAA ATATTTAGAA TCCGAAGTCC AAAGATCTTC TGCAATCTTG TCAAATCCTA
Group_IV-I   AAGAAATTAA ATATTTAGAA TCCGAAGTCC AAAGATCTTC TGCAATCTTG TCAAATCCTA
Group_IV-I   AAGAAATTAA ATATTTAGAA TCCGAAGTCC AAAGATCTTC TGCAATCTTG TCAAATCCTA
Group_IV-I   AAGAAATTAA ATATTTAGAA TCCGAAGTCC AAAGATCTTC TGCAATCTTG TCAAATCCTA
MycSu160_m   AAGAAATTAA ATATTTAGAA TCCGAAGTCC AAAGATCTTC TGCAATCTTG TCAAATCCTA

             ....|....| ....|....| ....|....| ....|....| ....|....| ....|....|
                 3005      3015      3025      3035      3045      3055
Group_IV-I   ATTTCTTAGC AAAAGCACCT AAAGAAAAAA TTGAATTAGA AAAATCTAAA TTAGAAGATT
Group_IV-I   ATTTCTTAGC AAAAGCACCT AAAGAAAAAA TTGAATTAGA AAAATCTAAA TTAGAAGATT
Group_IV-I   ATTTCTTAGC AAAAGCACCT AAAGAAAAAA TTGAATTAGA AAAATCTAAA TTAGAAGATT
Group_IV-I   ATTTCTTAGC AAAAGCACCT AAAGAAAAAA TTGAATTAGA AAAATCTAAA TTAGAAGATT
Group_IV-I   ATTTCTTAGC AAAAGCACCT AAAGAAAAAA TTGAATTAGA AAAATCTAAA TTAGAAGATT
```

```
MycSu160_m   ATTTCTTAGC AAAAGCACCT AAAGAAAAAA TTGAATTAGA AAAATCTAAA TTAGAAGATT

             ...

Group_IV-I   ATA
Group_IV-I   ATA
Group_IV-I   ATA
Group_IV-I   ATA
Group_IV-I   ATA
MycSu160_m   ATA
```
